# Combinatorial expression motifs in signaling pathways

**DOI:** 10.1101/2022.08.21.504714

**Authors:** Alejandro A. Granados, Nivedita Kanrar, Michael B. Elowitz

## Abstract

Cell-cell signaling pathways comprise sets of variant receptors that are expressed in different combinations in different cell types. This architecture allows one pathway to be used in a variety of configurations, which could provide distinct functional capabilities, such as responding to different ligand variants. While individual pathways have been well-studied, we have lacked a comprehensive understanding of what receptor combinations are expressed and how they are distributed across cell types. Here, combining data from multiple single-cell gene expression atlases, we analyzed the expression profiles of core signaling pathways, including TGF-β, Notch, Wnt, and Eph-ephrin, as well as non-signaling pathways. In many pathways, a limited set of receptor expression profiles are used recurrently in many distinct cell types. While some recurrent profiles are restricted to groups of closely related cells, others, which we term pathway expression motifs, reappear in distantly related cell types spanning diverse tissues and organs. Motif usage was generally uncorrelated between pathways, remained stable in a given cell type during aging, but could change in sudden punctuated transitions during development. These results suggest a mosaic view of pathway usage, in which the same core pathways can be active in many or most cell types, but operate in one of a handful of distinct modes.

## Introduction

In metazoans, a handful of core cell-cell communication pathways such as TGF-β, Notch, Eph-ephrin, and Wnt play critical roles in diverse developmental and physiological processes (Antebi, Nandagopal, et al., 2017; Gerhart, 1999; Li & Elowitz, 2019; Lim et al., 2015). Each of these pathways includes multiple, partly redundant, receptor variants that are expressed in distinct combinations in different cell types and interact in a many-to-many, or promiscuous, manner with corresponding sets of ligand variants (Figure 1A) (Derynck & Budi, 2019; Massagué, 2012; Okigawa et al., 2014; Rohani et al., 2014; Verkaar & Zaman, 2010; Wang et al., 2016). Within a given cell, the function of the pathway—which ligands it responds to, or which intracellular targets it activates—in general depends on which combination of components a cell expresses. For example, the TGF-β pathway, which plays pivotal roles in diverse developmental and physiological processes (David & Massagué, 2018), comprises 7 type I and 5 type II receptor subunits that combine to form heterotetrameric receptors composed of two type I and two type II subunits (Wrana et al., 1992). Cell types with distinct receptor expression profiles preferentially respond to distinct combinations of BMP ligands (Antebi, Linton, et al., 2017; Vilar et al., 2006), suggesting that different receptor combinations could provide distinct ligand specificities. Similarly, in mice, the Wnt pathway comprises a set of 10 Frizzled receptor variants that interact with 2 different LRP co-receptors, all of which are expressed in different combinations, and collectively control the cell’s response to combinations of Wnt ligand variants (Eubelen et al., 2018; Goentoro & Kirschner, 2009; Voloshanenko et al., 2017). The theme continues in the juxtacrine Notch and Eph-ephrin pathways where different membrane-bound ligand and receptor variants are expressed in diverse combinations and interact promiscuously to control which cells can signal to which others (Groot et al., 2014; Kania & Klein, 2016; Klein, 2012; Lafkas et al., 2015; LeBon et al., 2014; Sprinzak et al., 2010). Despite the prevalence of these promiscuous combinatorial architectures, it has generally remained unclear what pathway expression profiles exist and how they are distributed across cell types and tissues.

**Figure 1.**
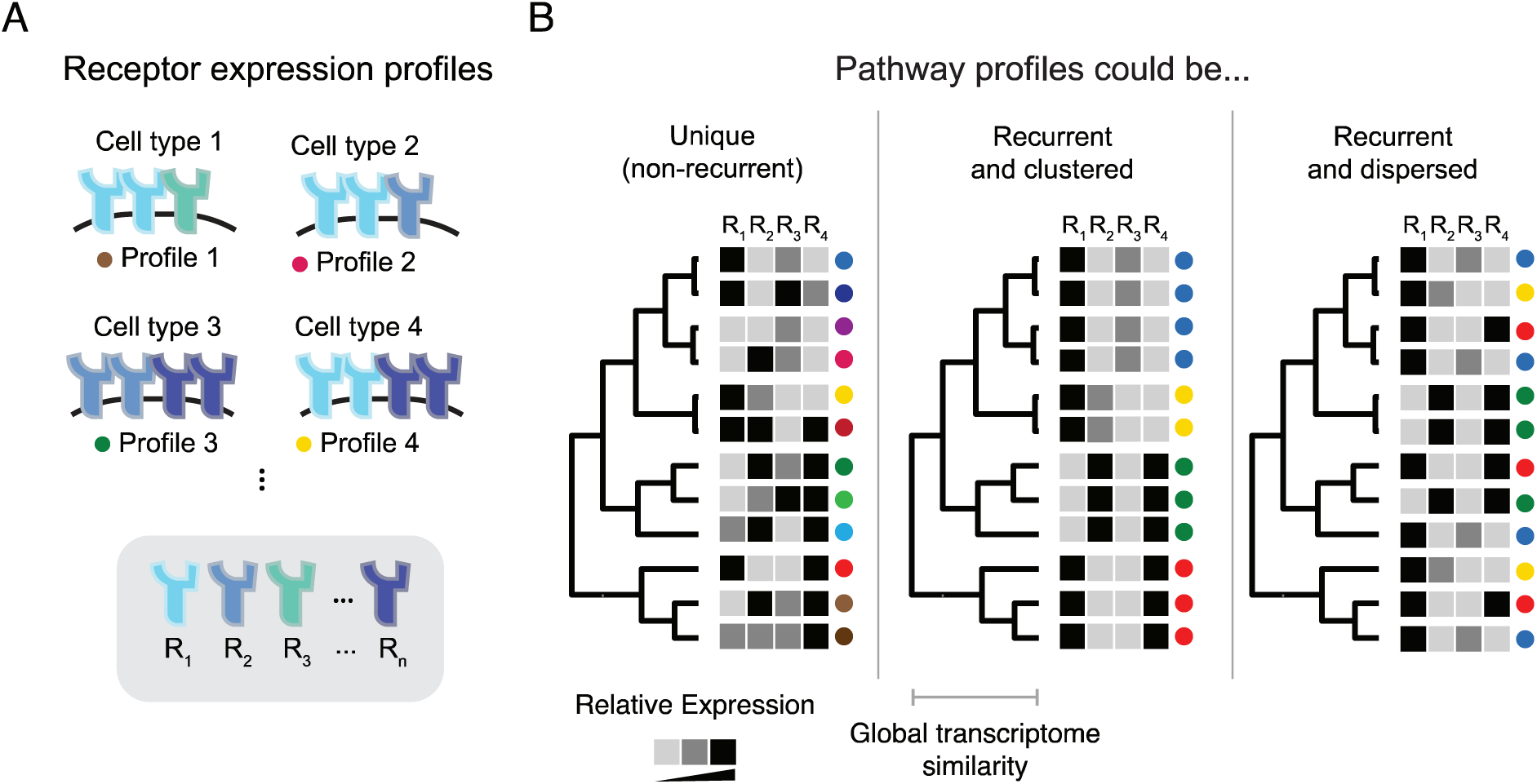
Pathway expression profiles could be distributed across cell types in different ways (schematic) A. Cell-cell signaling pathways comprise multiple variants of key components such as receptors (cartoons, Rn). These variants can be expressed in different combinations in different cell types. Colored dots identify receptor profiles for comparison with B. B. Cell types can be arranged hierarchically based on similarities among their global (genome-wide) gene expression profiles (dendrogram). A hypothetical signaling pathway profile for each cell type is indicated by the gray intensity in the corresponding row of squares. In principle, each cell type could have a unique signaling pathway profile (unique, left); exhibit a smaller set of recurrent profiles, each used by a set of related cell types (recurrent and clustered, middle); or exhibit signaling pathway profiles that recur even among otherwise distantly related cell types (recurrent and dispersed, right). These possibilities are not exclusive and it is possible that some pathways or subsets of cell types might operate in different regimes.

In principle, pathway expression profiles could be distributed across cell types in three qualitatively different ways. At one extreme, each cell type could express its own, completely unique, profile of pathway components (Figure 1B, left). In this case, one would observe as many distinct pathway profiles as cell types. Alternatively, sets of closely related (transcriptionally similar) cell types could share the same pathway expression profile (Figure 1B, center). This would result in fewer pathway profiles than cell types, and a correlation between the similarity of pathway profiles and the similarity of the overall transcriptomes of the cells in which they appear. Finally, a third possibility would be to observe a limited number of recurrent pathway profiles (as in the second case), but with individual profiles dispersed across multiple, distantly related cell types, rather than confined to sets of closely related cell types (Figure 1B, right). In this regime, otherwise similar cell types could exhibit divergent profiles for the pathway of interest, while, conversely, more distantly related cell types would converge on similar pathway profiles. In this last regime, a limited repertoire of profiles, which we term “pathway expression motifs,” are re-used in diverse cell contexts. Assuming that differences in pathway profile confer corresponding differences in ligand responsiveness or other properties, each of these regimes implies something different about the number and distribution of functionally distinct signaling modes for a pathway of interest.

Previously, systematically distinguishing among these potential classes of behavior would be difficult. Recently, however, single-cell RNA sequencing (scRNA-seq) cell atlases have begun to provide comprehensive gene expression profiles across most or all cell types in embryos and adult organisms. For example, the Tabula Muris project provided expression profiles for ∼100,000 cells across 20 organs in adult mice (Tabula Muris Consortium et al., 2018). This data set was later augmented with studies of mice at two additional ages (Tabula Muris Consortium, 2020). In parallel, scRNA-seq studies of embryonic development have similarly provided transcriptional profiles for the cell states in the early embryo (Grosswendt et al., 2020) and specific organs later in organogenesis (He et al., 2020). Collectively, these data provide an opportunity to determine the combinatorial structure of pathway expression.

Here, we introduce a statistical framework to identify pathway expression profiles and characterize their distribution across cell types in an aggregated data set spanning multiple atlases. This approach allowed us to identify the pathway expression motifs described above (Figure 1B, right) as well as “private” profiles that are limited to sets of closely related cell types (Figure 1B, middle) in core communication pathways including TGF-β, Notch, Eph-ephrin, and Wnt. These results suggest that each pathway can operate in a handful of distinct “modes.” Further, the mode used by one pathway appears to be independent of those used by other signaling pathways. Dynamically, pathway modes can remain remarkably stable during aging, or change suddenly as cells progressively differentiate during development. Together, these results provide a combinatorial view of signaling pathway states and suggest that many of the most central pathways can exist in a handful of different modes, which, in the future, may be studied independently of the cell types in which they appear.

## Results

### Integration of cell atlas data sets

To analyze pathway expression profiles across a broad diversity of cell types, we first compiled data from multiple adult and developmental cell atlas data sets (Figure 2A, Table 1). These included the Tabula Muris cell atlas (Tabula Muris Consortium et al., 2018), which comprises 40,000 cells distributed across 18 organs from a 3 month old mouse, as well as Tabula Senis (Tabula Muris Consortium, 2020), which augmented these data with ∼100,000 additional cells from mice aged 1, 18, 24, and 32 months. We also included two early developmental whole embryo atlases from E6.5 to E8.5 (Grosswendt et al., 2020; Pijuan-Sala et al., 2019a), and a forelimb organogenesis atlas from E10.5 to E15 (He et al., 2020). Each of these data sets also contained a cell type annotation for each cell based on expression of known markers. Altogether, the aggregated data set included expression profiles and cell type annotations for ∼700,000 individual cells.

**Figure 2.**
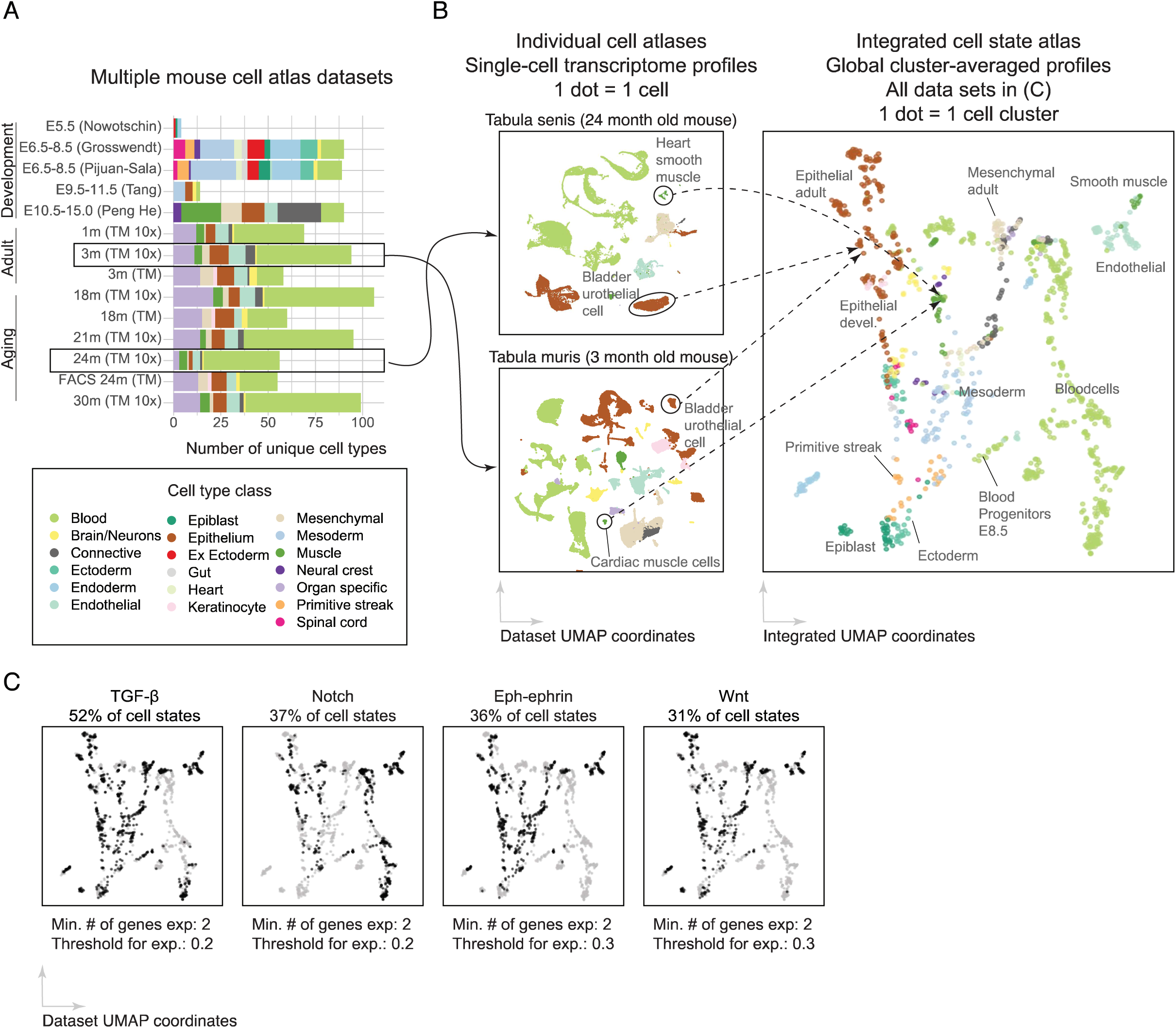
Integration of scRNA-seq atlas data reveals widespread expression of signaling pathway components. A. We integrated 14 published developmental and adult scRNA-seq datasets spanning different stages in the mouse lifespan from embryonic development to old age. These data sets differ in their representation of organs and cell type classes (colors). B. To generate an integrated cell state atlas, we first independently clustered each scRNA-seq dataset, treating distinct time-points in the data set separately (Methods). We then averaged expression over all cells in each cluster to yield a “cell state” profile for that cluster, and represented each cluster by a single dot in an integrated cell state atlas data set (UMAP on right). Colors are consistent with the legend in (A). Notably, this integration captures cell type similarity across different datasets and sequencing technologies. C. Components of core signaling pathways are broadly expressed. Black or gray dots show clusters whose pathway components are expressed above or below threshold, respectively.

**Table 1.**
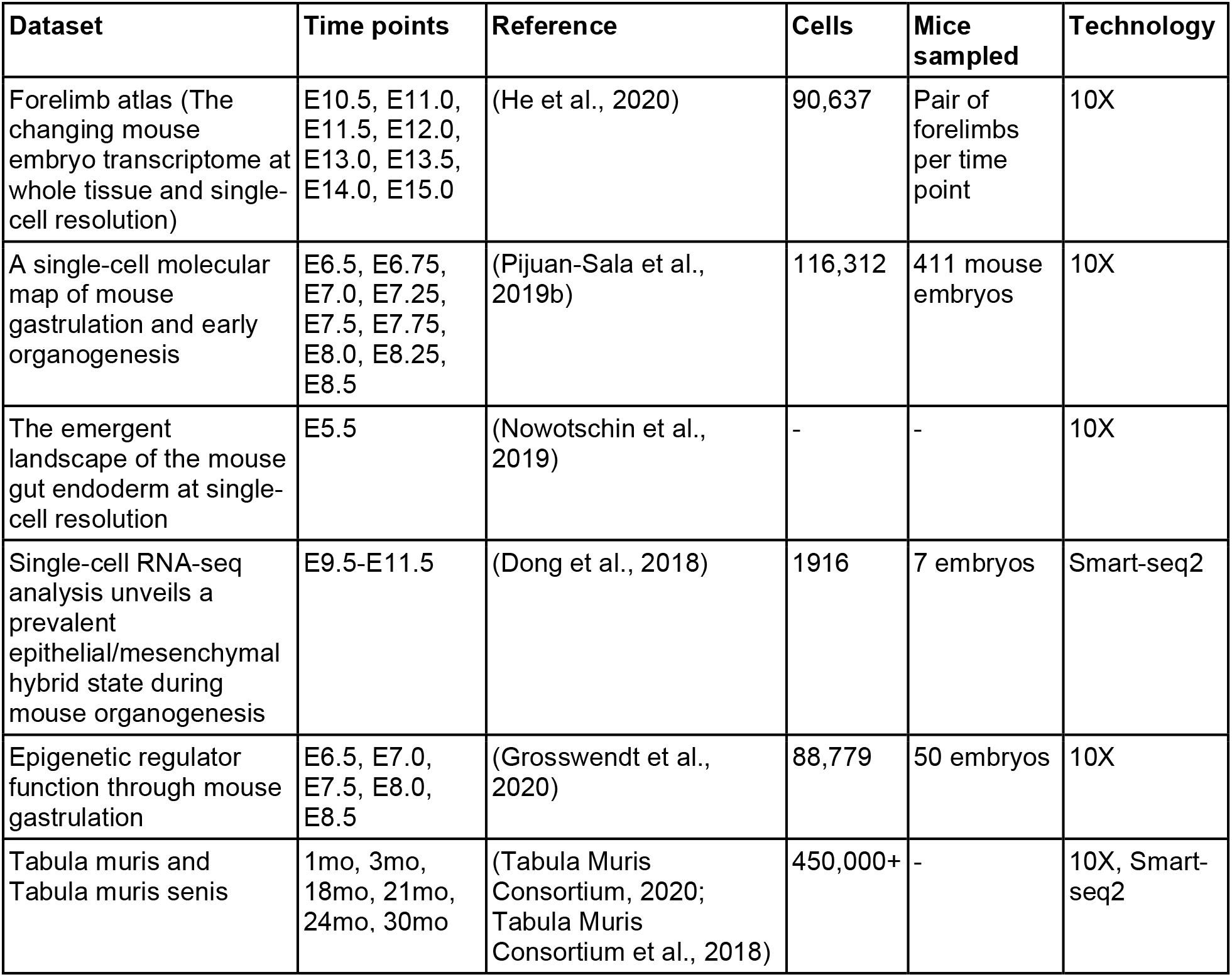
Single-cell data sets used in this work.

To allow a unified analysis of these data, we clustered the global transcriptional profiles from each dataset independently. This procedure resulted in 1206 clusters, spanning 917 unique cell type annotations (e.g. “Organ: Lung, cell type:endothelial, age: 3m”), providing a unified data set for further analysis (Figure 2B, Methods). For simplicity, in this work, we will refer to each global gene expression cluster as a “cell state” and not distinguish between formal “cell types” and other levels of variation. This clustering procedure and the cell states recovered from each dataset matched previous published analyses (Fig. 2–figure supplement 1A).

To focus on expression differences between cell states, reduce the complexity of the data set, and minimize the impact of measurement noise, we computed the average transcriptome profile of each one of the 1206 clusters (Methods), similar to other recent integration approaches (Qiu et al., 2021). Similar cell states in different data sets shared similar expression profiles, including for the specific pathways discussed below (Figure 2–figure supplement 1B). A UMAP projection displays the variety of cell classes comprising the integrated atlas (Figure 2B, right). We note that cluster averaging potentially eliminates biologically meaningful gene expression variability within a cluster. However, pairs of genes that were highly expressed within a cluster average also showed significant co-expression in single cells (p < 0.001; Figure 2–figure supplement 1C). The integrated, cluster-averaged dataset provides a basis for analyzing systematic changes in pathway gene expression between cell states in embryonic and adult contexts.

### TGF-β receptors exhibit recurrent expression profiles

Using the integrated data set, we first focused on the TGF-β pathway. A functional TGF-β pathway requires expression of at least one type I and one type II receptor subunit. Across the 1206 cell states, approximately half met this criteria, expressing at least one receptor of each type above a minimum threshold (Figure 2C, Methods). The most prevalent receptors, Bmpr1a and Acvr2a, were expressed in ∼10 times more cell types than the least prevalent, Acvr1c and Bmpr1b (Figure 3–figure Supplement 1A). Nearly every receptor subunit was co-expressed with each other receptor subunit in at least some cell types (Figure 3–figure supplement 1C). Even Acvrl1 and Bmpr1a, which were mainly expressed in endothelial and epithelial cells, respectively, were also co-expressed in mesenchymal cells (Figure 3–source data 1). Exceptions included Bmpr1b and Acvr1c, which were less prevalent overall and were co-expressed with a more limited set of other subunits (Figure 3–figure supplement 1C). Overall, these results provided TGF-β transcriptional expression profiles across cell types and revealed that they were strongly combinatorial.

To test whether certain receptor profiles recurred across cell types as in Figure 1B, middle and right panels, we clustered cell types based only on their TGF-β pathway expression profiles (Figure 3A). To detect recurrent profiles, we computed the silhouette score, which compares the separation of points between clusters to the separation of points within a cluster, and penalizes for both over- and under-clustering (Figure 3–figure supplement 2A) (Rousseeuw, 1987). The silhouette score provides a metric to quantify the approximate number of distinct clusters in a dataset. We compared the silhouette from actual profiles to those determined from randomized data sets in which the expression level of each receptor was independently scrambled among cell types (Figure 3–figure supplement 2B, black and gray lines). Subtracting the randomized silhouette score from that of the actual profile, and dividing by the standard deviation of randomized data, we obtained a z-score that quantifies how much the silhouette score from the actual profiles deviates that observed in the randomized control data, for a given number of clusters *k*. Finally, we selected the optimal number of clusters, *k_opt_*, that maximized this z-score (Figure 3—figure supplement 2B, blue). Altogether, this analysis revealed that 622 cell states expressing TGF-β receptors, collectively exhibit only about 30 distinct, recurrent pathway expression profiles (Figure 3A). Critically, every receptor subunit was expressed in at least one of these profiles, consistent with a combinatorial view of receptor utilization.

**Figure 3.**
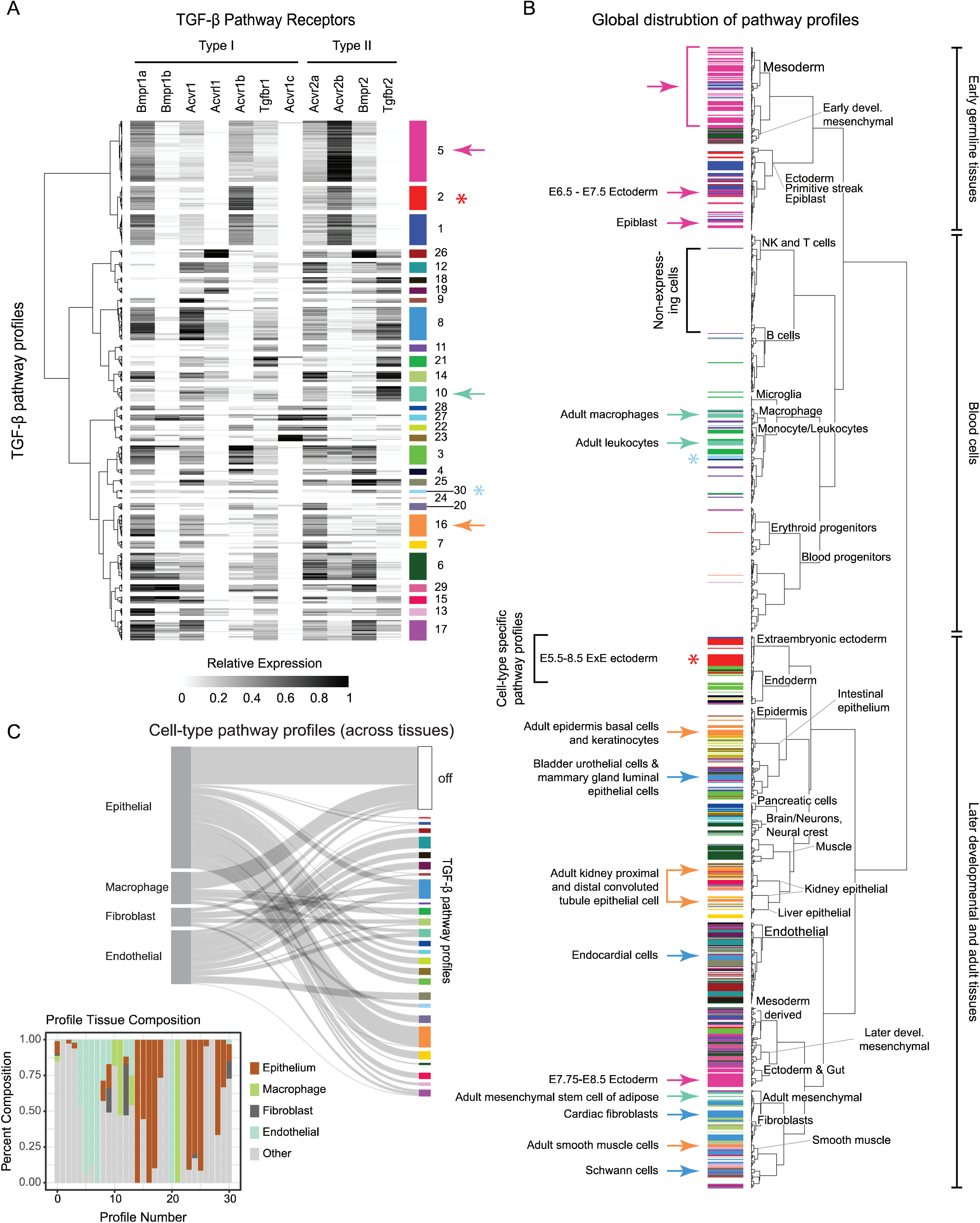
TGF-β receptors exhibit recurrent and dispersed pathway expression profiles. A. Silhouette score analysis (Figure 3-figure supplement 2A) identified approximately 30 TGF-β receptor expression profiles, indicated as color-labeled groups of rows. Colored arrows indicate examples of dispersed profiles highlighted on the global cell fate dendrogram in B. Asterisks indicate private profiles, also shown in B. Dendrogram at left represents similarity among different profiles. Each gene is standardized to a range of 0-1 across all cell types (grayscale). B. Distribution of TGF-β receptor expression profiles across cell types. The global cell type dendrogram was computed using a cosine distance metric applied to the integrated transcriptome data set in a 20-component PCA space constructed from 4,000 highly variable genes (HVGs). Arrows indicate featured TGF-β profiles that are broadly dispersed across cell types, while asterisks indicate examples of private profiles. Cell types that do not express TGF-β receptors have no color (white). Colors match those in A. Note that blood cell types are relatively lacking in expression of TGF-β receptors. C. Key cell type classes, including epithelial, macrophage, fibroblast, and endothelial cell types, each span multiple TGF-β profiles. The white bar (top right) indicates the non-expressing profile. Profiles are ordered to maximize the similarity of adjacent profiles. Each cell class mapped to multiple distinct pathway profiles, yet differed in their profile diversity. For example, epithelial cells comprise a broad spectrum of 18 distinct profiles, whereas macrophages and endothelial cells are primarily restricted to smaller subsets of more closely related profiles. Inset, cell type composition of each TGF-β profile, where “other” includes all cell states in the atlas that do not fall into the epithelial, macrophage, fibroblast or endothelial cell types.

### TGF-β pathway expression motifs appeared in diverse cell types

Having identified recurrent pathway expression profiles, we next asked how they were distributed across cell types, as in Figure 1B. To answer this question, we first visualized TGF-β pathway expression profiles on the dendrogram of global cell types (Figure 3B—supplementary file 1). We color-coded each profile in Figure 3A and then annotated each cell state on the global dendrogram with the color corresponding to its TGF-β profile (Figure 3B). Strikingly, many profiles were broadly distributed over diverse cell types (Figure 3B, colored arrows). For example, profile 10 (mint green) appeared in adult macrophages and leukocytes as well as mesenchymal adipose stem cells. On the other hand, a smaller number of pathway profiles showed the opposite behavior. They were restricted exclusively to a particular clade of closely related cell states (Figure 3B, colored asterisks). These results suggest that TGF-β could exhibit both pathway motifs and private profiles.

One potential explanation for the dispersion of recurrent pathway profiles could be if general classes of cell types, such as macrophages, fibroblasts, epithelial cells, or endothelial cells each adopted a particular, characteristic profile, irrespective of their tissue or organ context. For example, a pathway profile could appear dispersed if it occurred in a broad set of otherwise diverse macrophage cell types. We therefore used a Sankey diagram to visualize the relationship between each of these four cell type classes, based on cell type annotations in the atlas, and the full set of TGF-β profiles (Figure 3C). Some classes, such as epithelial cells, used more diverse TGF-β profiles than others, such as endothelial cells. Nevertheless, each of the four cell type classes mapped onto multiple TGF-β profiles. Conversely, most of the profiles appeared in multiple cell type classes or cell types (Figure 3C, inset). These results rule out these cell type classes as an explanation for dispersed use of recurrent pathway profiles, and suggest that pathway profile usage is based on other aspects of cell states.

To more systematically and quantitatively characterize the distribution of each pathway profile, we defined the “dispersion” of a given TGF-β profile as the mean value of the pairwise euclidean transcriptome distances among all cell types that express it, computed in the space of the 100 most significant principal components (Figure 4A). About 60% of TGF-β profiles were predominantly observed in specific sets of closely related cell types (Figure 4B, points between dashed lines). By contrast, 40% of TGF-β profiles were dispersed more broadly, often spanning distantly related cell types (Figure 4B, points above expected range). In fact, this subset of TGF-β profiles exhibited cell type dispersion levels approaching those expected if TGF-β profiles were assigned to cell types randomly (Figure 4C, blue versus black lines). Based on this analysis, we defined pathway expression motifs as profiles whose mean cell type dispersion exceeded a cutoff. For most analysis here, we set this cutoff at the 90th percentile of dispersions among groups of globally similar cell types (Figure 4D, Methods). Alternative dispersion metrics produced broadly similar, but not identical, motif sets, indicating some sensitivity to the definition of dispersion (Figure 4–figure supplement 1C). Finally, we note that this criteria is sensitive to an arbitrary threshold, the motif cutoff, here chosen at the 90th percentile. Reducing the motif cutoff would allow less dispersed profiles to be classified as motifs.

**Figure 4.**
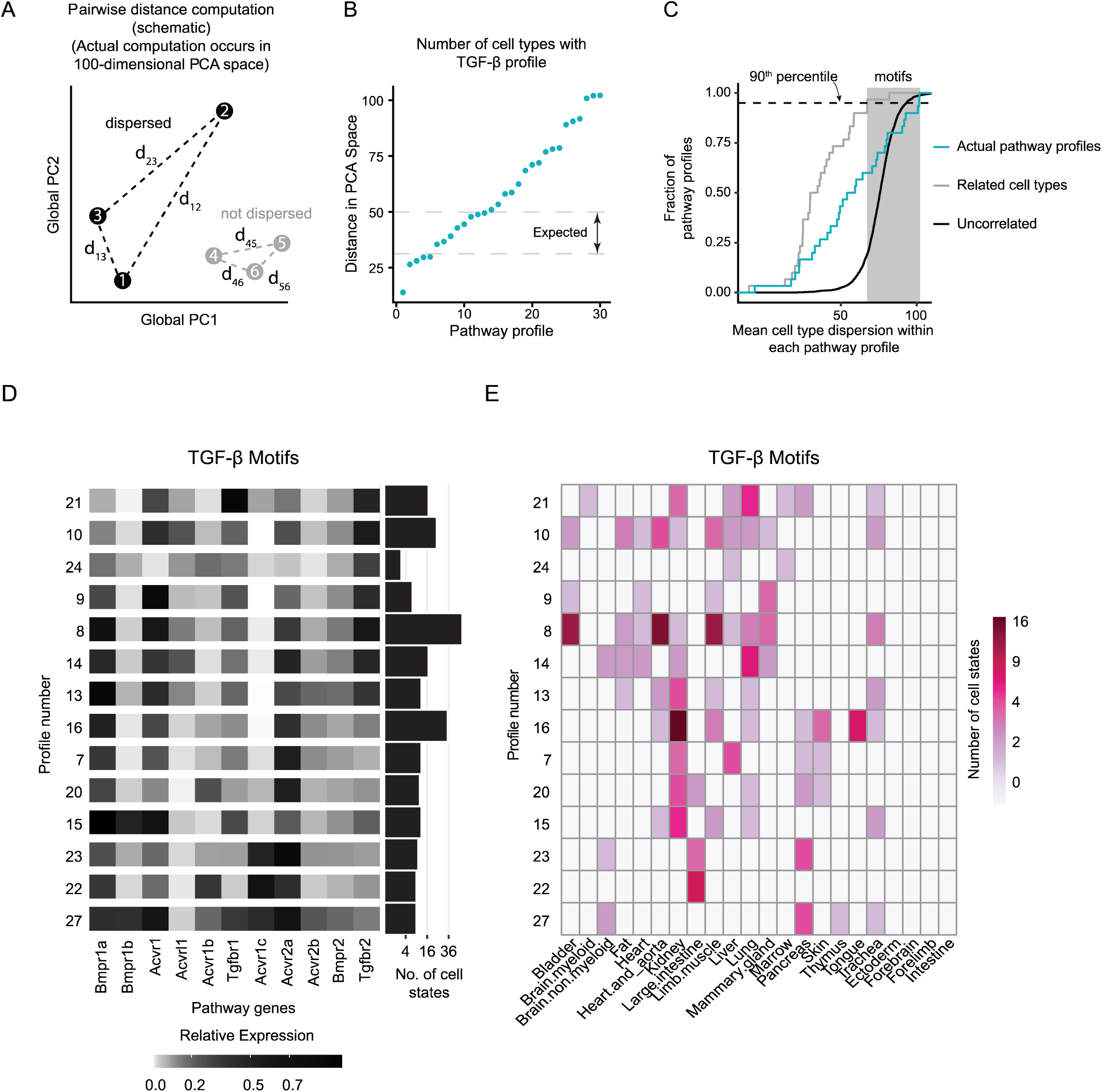
TGF-β expression motifs are dispersed across cell types and organs. A. We defined the dispersion of a receptor expression profile to be the within-class pairwise distance computed in a 100 dimensional PCA space constructed from the top 4,000 highly variable genes (HVGs) (left). Dispersed profiles (black) show high cell type diversity, whereas non-dispersed profiles (gray) are closer together in PCA space. B. The dispersion of actual TGF-β expression profiles. Dashed lines indicate the range of dispersions obtained for scrambled profiles. Note the large number of profiles with larger dispersions than expected from random profiles. C. Empirical cumulative distribution functions of TGF-β profile dispersion. The observed dispersion distribution (turquoise) lies between the extremes of cell type-specific profiles (gray) and profiles obtained by randomizing cell type distances by shuffling cell type labels (black). We classified motifs in the shaded region, defined as being in at least the 90th percentile of the related cell type dispersion distribution (gray) as motifs. D. We identified 14 TGF-β motifs, displayed in ranked order of dispersion from most (top) to least (bottom) dispersed. For each motif, the number of cell states in which it appears is indicated by the histogram at right. E. TGF-β motifs (rows) are broadly distributed across different tissues and organs (columns). Each matrix element represents the number of cell states in the indicated tissue or organ expressing the corresponding motif. Note that most motifs are expressed in multiple tissues or organs and most tissues or organs contain multiple motifs.

To better understand the structure of motifs, we also examined expression correlations among individual BMP receptors. Among cell states expressing pathway motifs, almost half of the receptor pairs (25/55) showed no significant correlation, with the remaining pairs exhibiting a mix of positive and negative pairwise correlations (Figure 4–figure supplement 1A). For example, Bmpr1a was positively correlated with Acvr1 and Acvr2a, while Acvrl1 and Tgfbr2 were strongly correlated, with Acvrl1 expressed in a subset of cell types that expressed Tgfbr2. Acvrl1 and Tgfbr2, which were previously shown to mediate signaling by BMP9, could also function together as a module in this context (Chen et al., 2013).

TGF-β pathway motifs exhibited several interesting features. First, they were enriched for expression of the type I receptors Bmpr1a and Acvr1, as well as the type II receptor Acvr2a. In fact, almost all motifs co-expressed all three of these receptor subunits (Figure 4D). On the other hand, Bmpr1b, Acvrl1 and Acvr1c were the least represented receptor subunits, appearing in only 3, 3, or 4 of the motifs, respectively. The most prevalent motif, 8, was expressed in 9 different mouse organs and is similar to the profile of NMuMG mammary epithelial cells, which were shown to compute complex responses to ligand combinations (Antebi, Linton, et al., 2017; Klumpe et al., 2020) (Figure 4D, rows). Motif 8 included the type 1 subunits Bmpr1a, Acvr1, and Tgfbr1, as well as the type II subunits Acvr2a, and Tgfbr2. Motif 15, which is similar to motif 8 but with more Bmpr1b, was shown to exhibit reduced complexity of combinatorial ligand responsiveness (Klumpe et al., 2020), suggesting that even a change in a single receptor between profiles could be functionally significant.

Motifs were broadly distributed across the organism, with some appearing in as many as 9 different mouse organs (Figure 4E, rows). Conversely, multiple motifs appeared in the same organ. For example, the adult kidney included cell states with 9 different TGF-β receptor expression motifs (Figure 4E, columns). These results underscore the breadth of the dispersion of the pathway motifs.

In contrast to motifs, other TGF-β profiles recurred in multiple cell types but exhibited low dispersion, as in Figure 1B, middle panel (Figure 4–figure supplement 1B). One of these groups, consisting of profiles 1,2, and 5, was in fact dispersed among diverse developmental cell types, including the primitive streak, ectoderm derivatives, and mesodermal tissues. However, it received a lower dispersion score due to the relative similarity of early embryonic cell types compared to adult cell types. We therefore classified these profiles as a developmental motif (Figure 3B, hot pink). These three profiles expressed a combination of Bmpr1a and Acvr2b, and resembled the BMP receptor profile previously identified in mouse embryonic stem cells, suggesting that the early embryonic receptor profile is stably maintained during early germ layer cell fate diversification (Klumpe et al., 2020).

By contrast, profiles 29 and 30 were each confined to a single set of closely related cell types: chondrocytes (E13.5-E15.0) and macrophages, respectively. Because they were tightly associated with a particular set of cell types, these profiles are effectively the opposite of a motif, and we refer to them as “private” profiles. Notably, these private profiles both expressed Bmpr2, which is less prevalent compared to other receptors. Nevertheless, Bmpr2 is not a marker of private profiles, as it is also expressed in dispersed motifs, such as motifs 8, 9, 10, 13, and 27 (Figure 4D). Together, these results suggest that the TGF-β pathway exhibits a set of recurrent and dispersed expression motifs, as well as a relatively small number of private profiles.

### Additional signaling pathways also exhibit pathway expression motifs

Other signaling pathways also exhibited recurrent expression profiles (Figure 5). Using the PathBank database of biological pathways (Wishart et al., 2020), we identified 56 different annotated biological pathways involved in signaling and other functions (Figure 5–source data 1). For each pathway, we assembled a corresponding list of genes, normalized their expression, clustered the resulting profiles, computed silhouette scores, and compared them to a null hypothesis in which the expression levels of each gene were independently and randomly reassigned to different cell types as described previously (Figure 5–figure supplement 1A). As with TGF-β, we identified the optimal number of clusters for each pathway by determining the peak value of the silhouette z-score.

**Figure 5:**
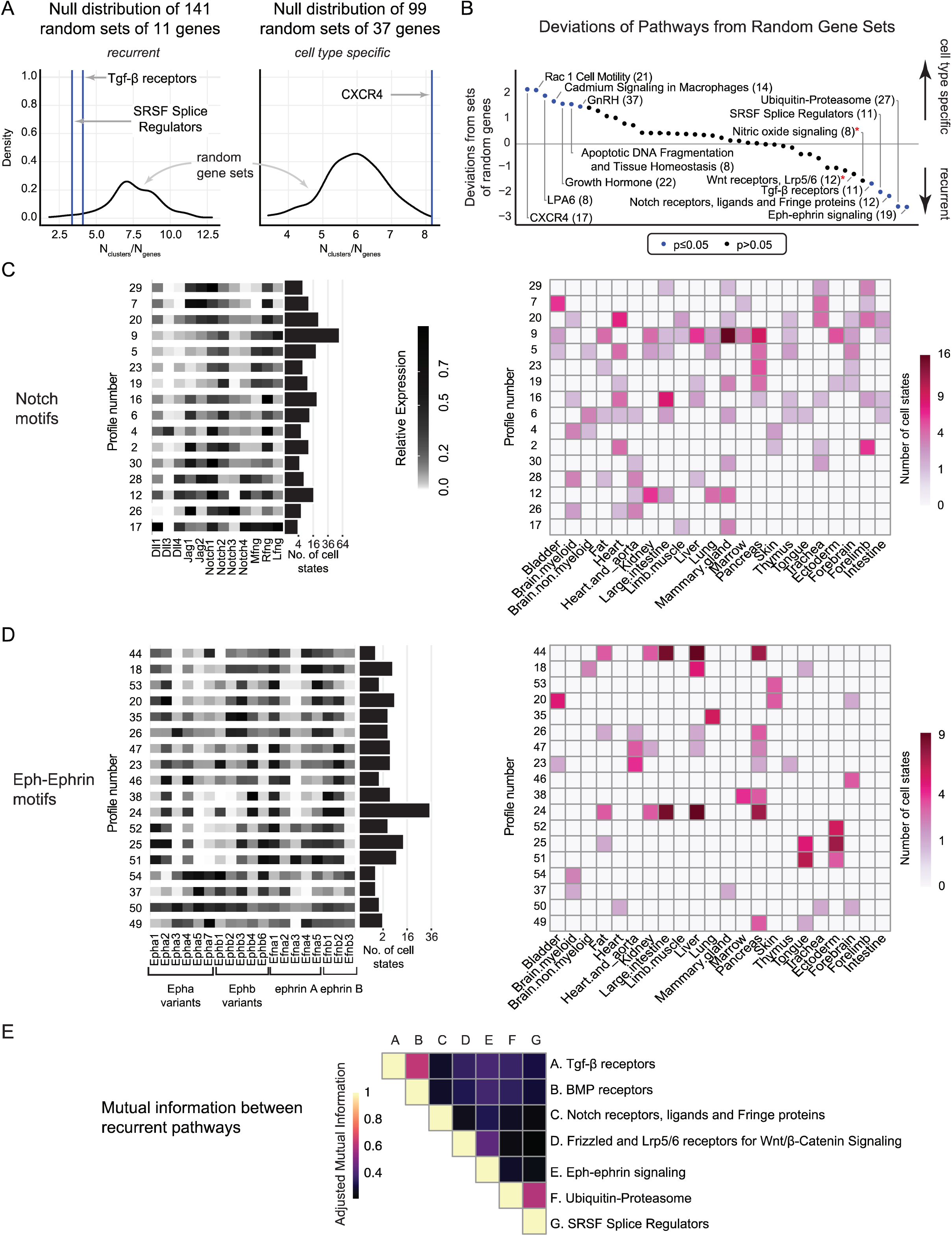
Expression motifs occur in multiple pathways. A. In order to classify a pathway as cell type-specific or recurrent, we compared the number of distinct profiles for a pathway (blue line) against a null distribution of the numbers of distinct profiles identified in random gene sets (black line). We computed these null distributions specific to the number of components in a pathway to avoid confounding the number of distinct profiles with pathway size, i.e., we would expect more combinatorial profiles for a pathway containing more genes. Left: examples of recurrent pathways (TGF-β and SRSF splice regulators), which have fewer clusters than expected from the null distribution. Right: example of pathway with more clusters than expected from the null distribution. B. Deviations of pathways from random gene sets. We curated 56 gene sets from the PathBank database and generated corresponding null distributions, analyzing each pathway for cell type-specific or recurrent behavior as in A. We normalized the number of identified clusters to the number of pathway components and computed the deviation of this ratio from the null distribution (y-axis). Negative deviations show that a signaling pathway has fewer clusters than expected for a given pathway size, indicating recurrence. By contrast, positive deviations occur when there are more clusters than expected, indicating strong cell type specificity. Pathways with significant deviations from the null distribution (adjusted p-value < 0.05) are highlighted in blue. Red asterisks indicate recurrent pathways that have strong, but not statistically significant, deviation from the null distribution. C. Motifs in the Notch pathway and their distribution across tissues and organs, similar to Figure 4D,E. D. Motifs in the Eph-ephrin pathway and their distribution across tissues and organs, similar to Figure 4D,E. E. Correlations in profile usage between pathways were quantified by the adjusted mutual information between their respective profile labels.

To classify pathways as recurrent or cell type-specific, we generated, for each pathway, a corresponding ensemble of ∼100 pseudo-pathways of the same size but composed of randomly selected genes (Figure 5A, black; Figure 5–figure supplement 1B, black). By clustering expression for each pseudo-pathway, we computed a null hypothesis distribution of *k_opt_* for each pathway of interest (Figure 5A, blue; Figure 5–figure supplement 1B, blue). We then calculated the difference between the observed number of clusters in the real pathway and the mean number of clusters found in the corresponding ensemble of pseudo-pathways (Figure 5B). Similar to TGF-β, several pathways exhibited fewer clusters than expected given their number of genes, indicating recurrent expression profiles (Figure 5B, right). These included core cell-cell communication pathways such as Notch, Ephrin, as well as the Srsf splicing protein family, including all 11 SR family splice regulatory proteins, and a protein degradation pathway defined at Pathbank consisting predominantly of different proteasome subunits (Wishart et al. 2020).

We also observed the opposite behavior: in some cases, pathway expression profiles in different cell states differed even more from one another than the expression levels of randomly chosen sets of genes. These pathways were thus the opposite of recurrent, or equivalently, highly cell type-specific, in their expression. They included CXCR4 (Figure 5A, right), Rac1, and Lysophosphatidic acid (LPA6) signaling. In each of these cases, the silhouette z-score exhibited no clearly defined peak and remained elevated compared to the null distribution, even as the number of clusters was increased (Figure 5B, left; Figure 5–figure supplement 1AB). Non-recurrent pathways may allow cells to fine tune a pathway to highly individualized requirements of each cell type. For example, in the CXCR4 or LPA6 pathways, this mechanism could allow each cell state to respond with a distinct amplitude and specificity to different sets of cytokines or LPA variants. These results indicate that some pathways have a non-recurrent structure dominated by private profiles.

Under our null hypothesis, signaling pathways were compared against a distribution of pseudo-pathways composed of randomly selected genes across the transcriptome (Figure 5B, *p*-values). We noted that this null distribution could underestimate the signal of the silhouette score since randomly selected genes exhibit different expression statistics compared to real pathways. Comparing against other randomized controls could increase the signal-to-noise for some pathways.

### Notch, Eph, and Wnt pathways exhibit dispersed expression motifs

#### Notch Signaling

We next asked whether other developmental signaling pathways similarly exhibited combinatorial expression patterns with recurrent, dispersed profiles. Based on their status as core signaling pathways and their recurrence scores (Figure 5B), we focused on Notch, Eph-ephrin, and Wnt.

In contrast to TGF-β and Wnt, which both use secreted ligands, the Notch pathway involves juxtacrine interactions between a set of membrane anchored ligands, including Dll1, Dll4, Jag1, Jag2, and the cis-inhibitor Dll3, and a set of four Notch receptors, Notch1-4 (Artavanis-Tsakonas et al., 1999; D’Souza et al., 2008; Siebel & Lendahl, 2017). Further, a set of three Fringe proteins (M-, R-, and L-Fng) modulates cis and trans ligand-receptor interaction strengths, both between adjacent cells (trans) as well as within the same cell (cis) (Kakuda et al., 2020; Kakuda & Haltiwanger, 2017). We therefore defined a minimal Notch pathway comprising 11 ligands, receptors, and Fringe proteins (Figure 5C). This definition excludes ADAM family metalloproteases, γ-secretase, the CSL complex, and other components, in order to focus specifically on ligands, receptors, and the Fringe proteins that directly modulate their interactions, all of which exist in multiple variants. We classified pathway expression as “on” if at least 2 of these genes were expressed above a minimum threshold of 20% of the maximum observed expression level across all cell types. With these criteria, the Notch pathway was “on” in 37% of cell states (450 out of 1200) (Figure 2C).

As with TGF-β, the Notch pathway exhibited combinations of co-expressed components, including receptors, ligands and Fringe proteins (Figure 5C). The pathway exhibited a peak Silhouette score at ∼31 cell clusters (Figure 5–figure supplement 1A), 16 of which qualified as motifs based on their dispersion scores (Figure 5-figure supplement 2, Figure 5C).

These profiles agreed with previous observations. For example, B cells (Notch motif 19) are known to express the Notch2 receptor and no ligands (Saito et al. 2003; Yoon et al. 2009). The combination of Notch1, Notch2 and Jag1 was prevalent, occurring in most of the motifs, which were distinguished by expression of other components (Figure 5C). Nevertheless, even among motifs expressing both Notch1 and Notch2, the ratio of the two receptors varied (compare Notch motifs 19 and 28, Figure 5C). Among the Fringe proteins, R-fng was expressed in all motifs, while L-fng and M-fng were restricted to a limited subset (Figure 5C). Nearly all motifs, with the exception of motif 26, which is expressed in cell types that comprise the blood vessels, co-expressed both ligands and receptors. Notch ligands and receptors are known to exhibit inhibitory (cis-inhibition) and activating (cis-activation) same-cell interactions that can generate complex interaction specifiities with other cell types expressing similar or different ligand and receptor combinations. The prevalence of multi-component Notch motifs could help explain complex Notch behaviors with the potential to send or receive signals to or from specific cell types (del Álamo et al., 2011; LeBon et al., 2014; Li & Elowitz, 2019; Nandagopal et al., 2019).

In addition to its expression motifs, Notch also exhibited a smaller set of ‘private’ expression profiles limited to closely related cell types (Figure 5–figure supplement 3A). Private motifs were used by muscle cells during forelimb development (profile 25), basal cells of the mammary gland (profile 21), mesodermal lineages at E7.0-E8.0, and the adult endothelium (profile 8). The private profiles exhibited greater expression of M-fng, and the Delta family ligands Dll1, 3, and 4 compared to the motifs (Figure 5–figure supplement 3A). Taken together, these results reveal that the Notch pathway uses a set of recurrent and dispersed combinatorial expression motifs, as well as private expression profiles in some lineages.

#### Eph-ephrin signaling

The most recurrent core signaling pathway in our panel was Eph-ephrin (Figure 5B, rightmost blue point), another juxtacrine signaling pathway that plays key roles in development, including tissue boundary formation, axon guidance, bone development, and vasculogenesis, among many other processes (Arthur & Gronthos, 2021; Cramer & Miko, 2016; Kania & Klein, 2016; Klein, 2012). Eph-ephrin signaling has also been implicated in numerous cancers (Astin et al., 2010; Merlos-Suárez & Batlle, 2008). The pathway implements juxtacrine communication bidirectionally between adjacent cells through combinations of Eph receptors and ephrin ligands, which are grouped into A and B families based on the specificity of their signaling interactions. Like Notch, Eph-ephrin interactions occur both in *cis* and in *trans*, and can also involve the formation of multi-component clusters (Dudanova & Klein, 2011). Furthermore, since the same ephrin ligand signaling through different Eph receptors can produce different and even opposite physiological responses (Seiradake et al., 2013), these features are consistent with the idea that component combinations could dictate signaling specificity.

Here, we tabulated the expression of 11 Eph variants and 8 ephrin variants, spanning both type A and B families (19 genes total). Silhouette analysis revealed a broad peak with a maximum at 54 clusters for the combined Eph-ephrin pathway (Figure 5–figure supplement 1A). Strikingly, all of these clusters exhibited co-expression of multiple Eph and ephrin variants (Figure 5D and Figure 5–figure supplements 2B, 3B). While Ephs and ephrins were generally not expressed in blood cell types (Figure 5–source data 2), they were broadly expressed in many others (Figure 5D). The Eph receptor expression profiles were also broadly distributed across these cell states, generating a set of motifs (Figure 5D). Inspection of the motifs revealed highly combinatorial expression patterns, co-expressing 3.67±1.88 and 2.89±1.23 Eph and ephrin variants, respectively, and nearly always expressing components from both A and B families. As with TGF-β and Notch, individual motifs often occurred in multiple organs and, conversely, individual organs often contained multiple motifs (Figure 5D, right). However, tissue coverage was more sparse than the other two pathways, possibly reflecting the greater number of distinct motifs (Figure 5D, left). These observed motifs agree with established signaling interactions observed *in vivo*. For example, an EphB4-EfnB2 signaling complex is known to regulate vasculature formation and maintenance in developing and adult mice (Salvucci & Tosato, 2012). Endothelial cells (motifs 24 and 47) notably co-expressed these components, in addition to other Eph receptor and ephrin ligand components.

The pathway also exhibited private profiles, which notably co-expressed a greater number of distinct components than the motifs (Figure 5–figure supplement 3B). Private profiles appeared in a variety of developmental tissues (profiles 17, 10, 7, 2, 1, and 14), as well as adult cell types (Figure 5–source data 2). Together, these results indicate that Eph-ephrin components are expressed in a combinatorial fashion with a mixture of motifs and private profiles, each broadly distributed across embryonic and adult tissues.

#### Wnt Signaling

Finally, as a fourth signaling pathway, we also analyzed Wnt, which plays critical roles in a vast range of developmental and physiological processes. Wnts can function as morphogens and are involved in regeneration, cancer, and disease (Grigoryan et al., 2008). Extracellular interactions between Wnt ligand and receptor variants exhibit promiscuity, with each ligand typically interacting with many receptor variants (Voloshanenko et al., 2017). Signaling involves Wnt ligands binding to Frizzled (Fzd1-10) receptors and low-density lipoprotein related co-receptors 5/6 (LRP5/6) to stabilize β-Catenin, allowing it to activate transcription of target genes (Goentoro & Kirschner, 2009; MacDonald & He, 2012; Mikels & Nusse, 2006). Wnt signaling has also been shown to have combinatorial features (Buckles et al., 2004).

The recurrence score for Wnt was slightly less than that of TGF-β and nitric oxide signaling (Figure 5B, red asterisks). Nonetheless, the pathway exhibited recurrent profiles. Silhouette score analysis showed a peak elevation at *k_opt_* = 30 profiles, similar to TGF-β, and was elevated compared to a null model of randomly scrambled pathways constructed from the same genes (Figure 5–figure supplement 1A). Strikingly, these profiles all exhibited co-expression of multiple Fzd variants, and all but two co-expressed both the Lrp5 and Lrp6 co-receptors (Figure 5–figure supplement 2C).

A subset of Wnt pathway expression profiles were broadly dispersed (Figure 5–figure supplement 3D). All of these high dispersion profiles co-expressed multiple Frizzled variants (Figure 5–figure supplement 3D). Conversely, most Frizzled variants were expressed in multiple high dispersion profiles. The exceptions were Fzd9 and Fzd10, which were expressed at much lower levels in most cell types, although Fzd9 was highly expressed in profile 28, along with other receptors (Figure 5–figure supplement 3C). These results show that the Wnt pathway also exhibits combinatorial expression motifs.

### Inter-pathway correlations reveal independent profile usage

Identifying combinatorial expression profiles in multiple pathways provokes the question of whether component configurations are correlated between pathways. For example, in the limit of tight coordination, cells expressing one TGF-β profile might always express a corresponding Notch profile. In the opposite limit, profiles from one pathway might be used independently of those from another pathway, suggesting a more mosaic cellular organization.

To quantify the correlation between expression profiles of different pathways, we computed the pairwise adjusted mutual information (AMI) between the profile labels of each pair of pathways across all cell types (numbers, Figures 2A, Figure 5–figure supplement 1A-C). The AMI metric quantifies the degree of statistical dependence between the two clusterings, controlling for correlations expected in a null, or completely independent, model. The full dataset of 1206 cell states was used for computing the pairwise AMI, assigning the profile label ‘0’ to cell states that do not express a given pathway. We visualized the results with a heatmap showing the pairwise AMI values across the main recurrent pathways (Figure 5E).

In general, most pathway-pathway correlations were weak (AMI < 0.4) (Figure 5E). To ensure that the AMI was indeed capable of capturing correlations, we included a subset of the TGF-β receptors (the 7 BMP receptors) as a separate pathway (“BMP receptors”). Given their overlapping components, TGF-β and BMP showed elevated AMI values of ∼0.6, as expected (Figure 5E). A notable exception was the strong correlation between the Ubiquitin-Proteasome pathway and SRSF splice regulators, which arose predominantly from developmental cell states expressing Ubiquitin-Proteasome profile 1 with SRSF profiles 1 and 2 (Figure 5–source data 2). Other pathway pairs, consisting of TGF-β, Wnt, or Eph-ephrin exhibited weaker relationships, whereas the Notch pathway showed little correlation with almost all other pathways. These results suggest that, at least for the limited set of components considered here, different pathways seem to adopt profiles largely independently of one another.

### Pathway profiles exhibit distinct dynamic behaviors during differentiation

The relative independence of profile selection between pathways provokes the dynamic question of when and how pathways switch profiles during development. At one extreme, profiles could switch in a stepwise fashion, changing one component at a time. At the opposite extreme, they could change multiple components simultaneously, directly switching from one profile to another. Further, either type of change could occur gradually or suddenly, and could be temporally synchronized or unsynchronized between different pathways.

Neural crest differentiation provides a well-characterized developmental process to address these questions. The neural crest is responsible for diverse cell types, including sensory neurons, autonomic cell types, and mesenchymal stem cells (Kléber et al., 2005; Simões-Costa & Bronner, 2015). Further, TGF-β, Notch, Eph-ephrin, and Wnt, all play key roles in its differentiation (Bhatt et al., 2013).

Soldatov et al. performed deep scRNA-seq analysis of neural crest development from embryonic day 9.5 cells using SMART-seq2 (Soldatov et al., 2019). We used the Slingshot package (Street et al., 2018) to construct pseudotime trajectories from these data and further identified 7 distinct pseudotime stages (Figure 6A). All expression counts were scaled to match the normalization used in the integrated atlas (Figure 2, Methods). This reconstruction recapitulated known cell fate trajectories, with neural crest progenitors differentiating into sensory neurons, autonomic neurons, and mesenchymal cells (Figure 6A). Except for a transient upregulation of Bmpr1b early on, the TGF-β profile was remarkably stable during the trajectory from progenitors to more differentiated cell types. The profile was dominated by Bmpr1a, Tgfbr1, Acvr2a, and Acvr2b (Figure 6B, first panel), closely matching profile 6 (Figure 3A), which occurs in the developing forebrain and spinal cord, adult mesenchymal, and adult podocyte cell types. This profile is potentially functional, as TGF-β pathway inhibition in neural crest stem cells leads to cardiovascular defects (Wurdak, 2005). These results indicate that a developmental pathway can retain a stable profile along a differentiation trajectory.

**Figure 6.**
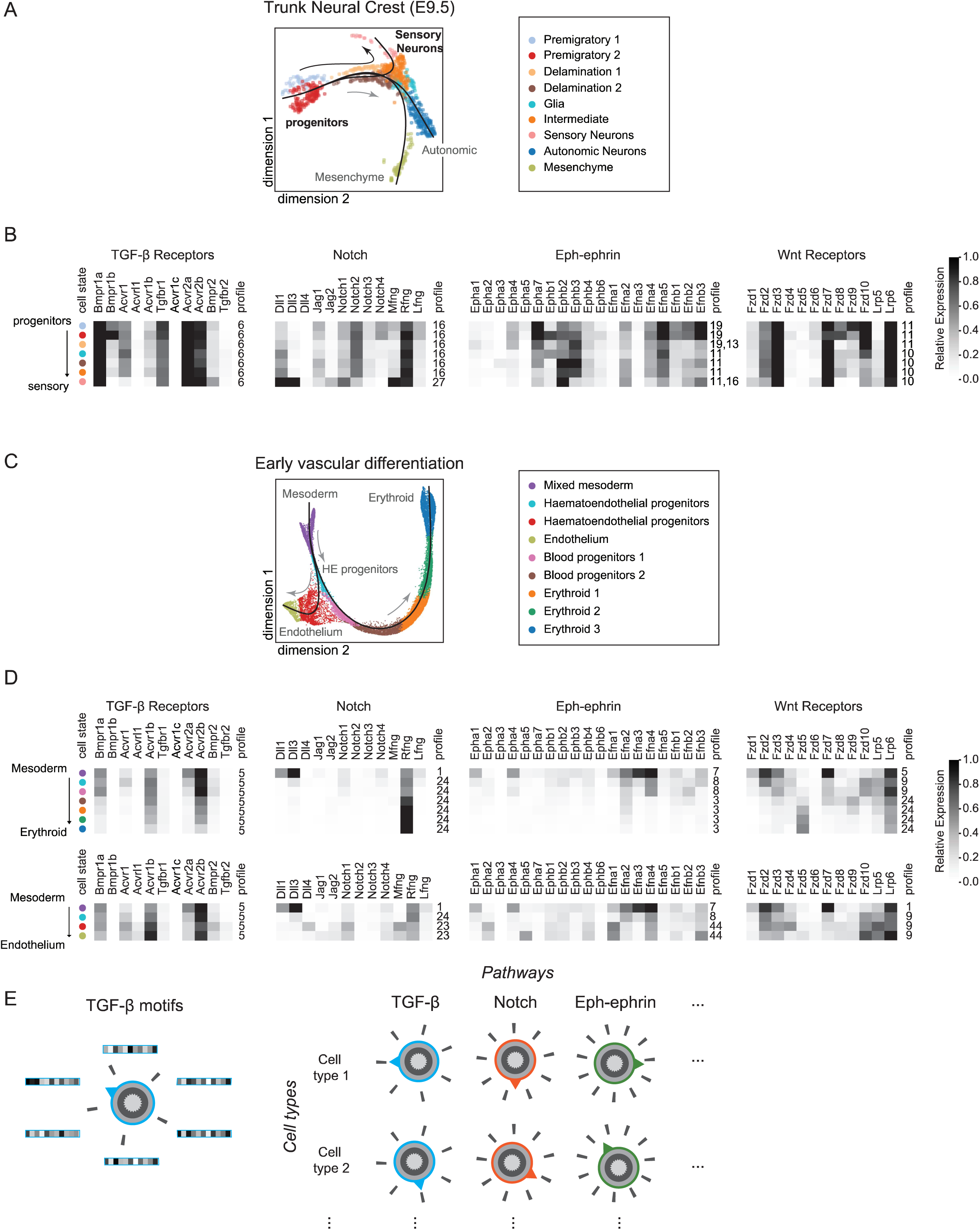
Developmental transitions of pathway profiles. A. Pseudotime trajectory analysis of the trunk neural crest (Soldatov et al., 2019) captures delamination of progenitors into three distinct cell fates in a ForceAtlas projection: sensory neurons, autonomic neurons, and the mesenchyme. Here, we follow the sensory neuron trajectory (black arrow). B. Developmental pathways show distinct expression dynamics in neural crest differentiation. For each pathway, corresponding mean expression profiles are shown in grayscale for each of the cell states indicated in A. Colored dots indicate which populations are being averaged. Profile numbers indicate the closest match to one of the reference pathway profiles shown in Figures 3A and 5-figure supplement 2. Two numbers are indicated for profiles that are approximately equally similar to the corresponding reference profiles. C. In early vascular differentiation (Pijuan-Sala et al., 2019b), mesodermal progenitors differentiate into endothelial and erythroid cell fates (gray arrows in ForceAtlas projection). D. Dynamics of four core pathways for each of the two trajectories in C: erythroid differentiation (upper row of heat maps) and endothelial differentiation (lower row). Colored dots indicate cell populations in C. Profile numbers indicate closest matches in reference profiles (Figure 3A, Figure 5-figure supplement 2). E. Mosaic view of profile usage (schematic). Cell states can express each of their pathways, using any of the distinct available profiles (indicated schematically by profile ticks). In this way, cell states can be thought of, in part, as mosaics built from combinations of available pathway profiles.

In contrast to the stability of TGF-β along this trajectory, Notch components exhibited a step-like transition at the end of the pseudotime trajectory (Figure 6B, second panel). Progenitors predominantly express the receptors Notch1 and Notch2; the ligands Dll1 and Jag1; and high levels of Rfng. This profile resembles Notch motif 16 (Figure 5C). Upon differentiation into sensory neurons, they switch on expression of Notch1, Dll3, and Mfng, as well as a lower level of Jag2, while down regulating Notch2, thus changing to private profile 27 (Figure 5C). Consistent with this analysis, profile 27 was independently derived from neural crest cells in the integrated data set (Figure 5–source data 2). A similar pattern of discrete change also occurred in the Wnt pathway, where expression shifted in ∼2 steps from profile 11 to profile 10 (Figure 6B, fourth panel). Thus, the transition to the sensory neural fate involves an abrupt multi-gene alteration of Notch and Wnt pathway components, neither of which was synchronized with changes in TGF-β.

By contrast, the dynamics of the Eph-ephrin pathway were more complex and gradual, with changes occurring in the expression of individual receptors at nearly every pseudotime stage. Eph-ephrin expression initially resembled profile 19 (Figure 5-figure supplement 2B), then switched more gradually to profile 11, before diverging slightly from it in the last pseudotime point (Figure 6B, third panel). Collectively, these results show that during neural crest development, different pathways can exhibit both stability and multi-step changes in their expression profiles.

As a second case, we analyzed hematopoiesis, which occurs in temporally and spatially overlapping waves in close proximity to blood vascular endothelial cells (Canu & Ruhrberg, 2021). Mesodermal hematoendothelial progenitors differentiate into both endothelium and erythroid cells (E7.5-E8.5), allowing analysis of how pathway profiles change during a branched differentiation trajectory (Figure 6C). Endothelial cells exhibit ‘private’ TGF-β profiles, characterized by expression of ACVRL1. Thus, this process provides an opportunity to analyze how pathway profiles change during a branched transition and how private profiles are acquired dynamically.

We clustered the subset of haemato-endothelial lineages from (Pijuan-Sala et al., 2019b) (15,645 single-cells), applied Slingshot to reconstruct branching pseudotime trajectories (Figure 6C), and then analyzed changes in TGF-β receptor expression profiles over these trajectories. In contrast to its stability during neural crest differentiation, the TGF-β profiles exhibited complex, dynamic changes during vascular differentiation. Mesodermal cells predominantly express Bmpr1a, Acvrl1, Tgfbr1, Acvr2a and Acvr2b, and Acvr2b, resembling profile 5, which is prevalent in early development (Figure 3A). Along the erythroid lineage, cells exhibited a gradual reduction in expression of all TGF-β receptors. Similar decreases in expression were also observed for receptors and ligands in other pathways (Figure 6D, upper row), and may reflect preparations for the dramatic events of erythropoiesis. By contrast, cells differentiating into endothelial fates maintained Bmpr1a and Acvr2b expression and additionally up-regulated Acvrl1, an endothelial-specific BMP receptor known to mediate signaling by BMP9 and BMP10, and required for angiogenesis (Tual-Chalot et al., 2014). Thus, while one lineage gradually turns off receptor expression, the other activates a distinct endothelial specific receptor profile. Looking more broadly at the four pathways during differentiation to endothelium, we see similar themes as observed in the neural crest differentiation: unsynchronized transitions to different profiles in different pathways. Together, these results show how pathways discretely and independently alter their expression profiles during different developmental lineages.

## Discussion

In multicellular organisms, a core set of molecular signaling pathways mediate a huge variety of developmental and physiological events. How can such a limited set of pathways play such a broad range of different roles? At a coarse level, each pathway may be considered competent for signaling in a given cell type if its receptors and other components are expressed and not inhibited by other cellular components. However, examining pathway expression patterns globally, as we did here, reveals a more subtle situation, in which pathways can be expressed in a finite number of distinct configurations, characterized by different expression levels for its components, all potentially competent to signal in response to suitable inputs. Each configuration could be functional in some contexts but nevertheless differ from other configurations in the specific input ligands it senses, or the downstream effectors it activates within the cell (Antebi, Nandagopal, et al., 2017; Buckles et al., 2004; Klumpe et al., 2020; LeBon et al., 2014; Li & Elowitz, 2019; Rohani et al., 2014; Su et al., 2020; Verkaar & Zaman, 2010).

To find out what configurations exist, we focused on cell-cell signaling pathways known to use sets of partially redundant component variants. Each of these pathways was already known to adopt multiple expression configurations in specific biological contexts. However, cell atlas data permit a systematic analysis of expression profiles in a broad set of cell and tissue contexts (Figures 2-5), revealing what pathway profiles are expressed, how they correlate with one another between pathways (Figure 5G), and how they change dynamically during aging and development (Figure 6).

The expression profiles of pathways are strikingly combinatorial. Across each of the four major pathways studied here, no two components exhibited identical expression patterns, and all were differentially regulated in some cell types. Further, almost all motifs comprised multiple receptor and/or ligand variants. The number of distinct expression profiles for each pathway was much smaller than one would expect if individual components varied independently. For instance, the Eph-ephrin pathway with 19 components exhibits ∼54 profiles, which is less than two-fold greater than the ∼30 profiles observed for the 11 TGF-β receptors, and far less than the 2^19^=524,288 pathway profiles one would expect if each of its 19 genes could independently vary between low and high expression states. Assuming that the pathway profile plays a key role in controlling pathway function, this finding suggests that analysis of a limited number of profiles could potentially explain pathway behavior in a much larger number of cell types.

Expression profiles for different pathways appeared to vary independently across cell types (Figure 5G). This observation argues against tight coupling of specific expression receptor profiles in one pathway with those in another. However, it does not rule out the possibility that signaling through combinations of pathways could play special roles in some cases (Muñoz Descalzo & Martinez Arias, 2012). Analysis of pseudotime trajectories also revealed that different pathways sometimes switch among motifs in a punctuated manner, and largely independently of one another. While we focused on the pathways that show strong motif signatures, it is equally important to note that other pathways predominantly used cell type specific, or private, profiles (Figure 5B), and even the pathways that we focused on here also contained some private profiles. Nevertheless, these results suggest a “mosaic” view of cells, in which each cell type adopts a particular motif or private profile for each of its general purpose pathways (Figure 6E).

Why use motifs? Motifs could provide a rich but limited repertoire of distinct functional behaviors for each pathway (Su et al., 2020). One appealing possibility is that each motif has a distinct but related signaling function that is retained in some way even in different cell types or contexts. For example, in a “combinatorial addressing” system, different ligand combinations could selectively activate sets of cell types based on their receptor expression profiles, to achieve greater cell type specificity in signaling (Klumpe et al., 2020; Su et al., 2020). A similar principle could apply to juxtacrine signaling pathways such as Notch and Eph-ephrin, where the combination of components expressed in a given cell type could control which other cell types it can communicate with, based on their own pathway expression profiles. To test this possibility, it will be important to determine what inputs each motif can respond to, and whether that specificity is retained across different cell contexts.

Several limitations apply to the findings reported here. First, pathway definition starts with a human-curated list of receptors, ligands, or other components or previously annotated pathway definitions. Different pathway definitions could potentially alter these results. Second, while comprehensive, the data sets used here are likely incomplete, and could miss profiles used only by rare cell types or could inaccurately report expression levels for weakly expressed genes.

Third, clustering is an imperfect representation of expression variation, potentially averages over subtle quantitative differences in individual component levels between cells. In particular, unsynchronized single cell dynamics, such as those that occur during Notch-dependent fate determination (Kageyama et al., 2018), could therefore be missed. Moreover, we explored signaling dynamics in only a few developmental trajectories. A broader exploration of more developmental processes could potentially reveal other types of dynamic behaviors beyond those shown here. Finally, subcellular localization patterns, post-translational modifications, alternative splice forms, and other types of regulation could diversify the functional modes of the pathway beyond what can be detected by scRNA-seq. However, as single cell technology continues to improve and expand to the protein level, we anticipate that it should be possible to obtain more precise views of pathway states.

The combinatorial nature of pathways makes it infeasible to experimentally characterize all possible configurations. Fortunately, however, a handful of motifs account for a large fraction of cell types, potentially enabling one to understand most of the functional repertoire of a pathway from a limited number of motifs and private profiles. While we focused on signaling here, the approach could be applied more generally to non-signaling pathways, such as splice regulation or protein degradation (Figure 5B). In the future, we anticipate that a functional understanding of pathway motifs could enable one to predict and control the activities of pathways in cell types based on their expression profiles.

## Acknowledgements

We would like to thank Heidi Klumpe, Rachael Kuintzle, Matthew Langley, James Linton, Benjamin Emert, Nicolas Pelaez-Restrapo, and other members of the Elowitz lab for suggestions and critical feedback on this work, as well as critical feedback from Matt Thomson, Kai Zinn, Yaron Antebi, and Miri Adler. This research was supported by the Allen Discovery Center program under Award No. UWSC10142, a Paul G. Allen Frontiers Group advised program of the Paul G. Allen Family Foundation, and by the National Institutes of Health grant R01 HD075335A. N.K. was supported in the summer of 2020 by the Samuel N. Vodopia and Carol J. Hasson SURF Endowment.

## Methods

### Clustering single cells and defining cell states

We obtained raw scRNA-seq matrices directly from the GEO repositories or specific locations indicated by the authors for the data sets appearing in the table below. Clustering of single cells started from the count matrices of single cells vs genes. First, we applied quality control (when needed, since some datasets were already filtered) by filtering out cells with high mitochondrial RNA content, low number of detected transcripts or low number of detected counts. We then applied a standard pipeline for clustering scRNA-seq data. Briefly, we applied principal component analysis and used the first 50 principal components as input for graph-based (Leiden) clustering using Scanpy (Traag et al., 2019; Wolf et al., 2018). Finally, we labeled the resulting clusters using the cell type annotations provided by the authors. All datasets analyzed in this study included ground truth cell type annotations that we use throughout the manuscript. All raw and processed data, along with scripts, are available at . Code can be found at https://github.com/nkanrar/motifs.git.

### Integration of multiple datasets

To integrate the above datasets into a single matrix of gene expression, we first generated a pseudo-bulk expression matrix for each dataset by averaging the log-normalized gene expression values of individual cells in a cluster. The resulting matrix has dimensions N x M, where N is the number of cell states in the dataset and M is the number of distinct genes. To account for differences in gene detection across datasets, we found the intersection of detected genes across all datasets and subsampled each matrix to include only genes that appeared in all data sets. The intersection of detected genes across all datasets comprised ∼11,000 genes that we then used for all downstream analysis. Having defined the intersection gene set, we concatenated individual datasets into a global average expression matrix containing 1206 clusters and ∼11,000 genes.

To normalize gene expression values from different datasets to a common scale, we applied a second round of normalization to the global expression matrix. First, we transformed the log-normalized matrix 𝛭 using the exponential function to obtain a matrix *M_ij_* of “counts” per gene: 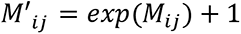. We then normalized, scaled and clustered the resulting matrix following the standard methods from Seurat v3 (total RNA counts per cell state = 1e4, 4,000 highly-variable genes and 50 principal components), which resulted in the clustering and UMAP shown in Figure 2. We verified that cell states from different datasets and sequencing technologies clustered together (Figure 2B), as an indication that the integrated and normalized UMAP recovers the biological diversity across development, adult and aging datasets.

### Clustering pathway expression profiles across cell states

All downstream analysis on pathway genes starts from the normalized pseudo-bulk gene expression matrix described above. We noticed that pathway genes showed different dynamic ranges in their expression across cell states. To give each pathway gene equal weight during clustering of pathway profiles, we applied a MinMax scaling for each gene, using the 95% percentile observed across all 1206 cell states as the maximum value. After scaling, each gene in the pathway had a dynamic range from 0 to 1, corresponding to the range of 0-95% of the maximum value in the data set for that gene. For each cell state, we classified a pathway as being “on” if at least two of the pathway genes showed expression above a threshold of 0.2 on this scale, meaning that the gene is expressed at a level of at least 20% of its maximum observed value. This threshold allowed us to filter out cell states in which all genes in the pathway are zero or showed low expression compared to most other cell states, and focus instead on the cell states showing combinatorial expression of multiple genes (Figure 2–figure supplement 1B, C). We computed all pairwise cosine distances between cell states with an “on” pathway profile, considering only the pathway genes, and applied hierarchical clustering to the resulting distance matrix (Figure 3A).

For each pathway, we found the approximate optimal number of clusters, *k*_opt_, using the silhouette score metric. After applying hierarchical clustering to the pathway expression matrix, one can define a number of clusters, *k*, by setting a depth cut-off and splitting the associated dendrogram (Figure 3A). We therefore computed the average silhouette score for a range of *k* values (from 3 to 100). To account for potential clustering artifacts, we randomized the pathway gene expression matrix, shuffling the expression values for each gene across cell states, and repeated the clustering procedure. By independently randomizing the matrix 200 times, we generated a null distribution for the expected silhouette score at different values of *k* (Figure 3- figure supplement 2B gray). From this null distribution, we computed z-scores for the silhouette scores obtained from the real pathway expression matrix and defined the optimal number of clusters, *k*_opt_, as the value of *k* with the most significant z-score (Figure 3—figure supplement 2B, dotted line).

### Defining motifs and private profiles based on cell type diversity

Having defined the *k*_opt_ clusters, or pathway profiles, we computed the diversity of cell states expressing each profile based on their transcriptome similarity. In principle, pathway profiles might comprise similar cell types (high transcriptome similarity) or sets of diverse cell types (low transcriptome similarity). We calculated their pairwise Euclidean distances in the PCA projection constructed from the top 4000 highly variable genes (50 principal components) to measure transcriptome similarity in a subset of cell states. We first verified that this metric was low for closely related cell states (as defined by their cell type annotation) and largest for randomly selected cell states (Figure 2—figure supplement 1B). We then defined *dispersion* as the average pairwise PCA distance among a subset of cell states.

To find the lower bound of dispersion, we computed the expected dispersion for related cell states by clustering their transcriptomes using the first 50 principal components, resulting in a global dendrogram of cell states (Figure 3B). We then identified the clustering threshold for the global dendrogram to obtain the same number of clusters *k* as observed for the pathway in question, therefore generating k groups of cell states that are each closely related. We then compared the distribution of dispersions for clusters of related cell states and the dispersions for cell states within the pathway profiles (Figure 4C). The dispersion distribution observed for related cell states (gray line Figure 4C) defines an approximate lower bound for the dispersion (Figure 4C). Conversely, we also computed dispersion values for randomly selected groups of cell states (Figure 4C, black). Random groups of cell states provide the dispersion expected if pathway expression states were completely uncorrelated with the overall expression similarity of the cells in which they appear. Finally, we defined a pathway profile as a *motif* if the cell states expressing it showed dispersion values higher than the 90% percentile value expected for related cell states (Figure 4C—shaded area). The 90% percentile threshold in dispersion identified pathway profiles expressed in the most diverse set of cell states. However, we observed additional pathway states that appeared dispersed among cell types but did not meet pass the 90% threshold. Therefore, this method could underestimate the number of dispersed pathway profiles and the threshold can be adjusted to allow a more flexible definition of pathway motifs.

In contrast to pathway motifs, “private” profiles are cell-state specific, effectively the opposite of motifs. By definition, private profiles are confined to sets of similar cell states and therefore show low dispersion values. To classify private profiles, we identified those profiles whose cell state dispersion overlapped with the expectation for highly-related cell states. Specifically, we considered profiles with dispersion < 30% percentile of the lower-bound distribution as “private.” For a pathway to be cell-state specific we expected the dispersion to be similar to that observed in closely related cell states. The threshold can be increased to allow for identification of other pathway profiles with dispersion values comparable to related cell states.

### Recurrence screening in multiple pathways

We calculated recurrence across multiple signaling and protein pathways from the PathBank (Wishart et al. 2020) database. First, we generated pathway expression matrices for 56 pathways annotated as ‘Signaling’ or ‘Protein’ in PathBank, excluding pathways with less than 7 genes. Next, we generated 200 pseudo-pathway expression matrices with the exact dimensions for each pathway expression matrix by randomly sampling genes from the transcriptome. We then generated a null distribution for the expected number of clusters in a typical set of genes in the transcriptome (Figure 5A) by following the procedure described above. Some pathways, however, did not show a clear peak in the z-score (Figure 5—figure supplement 1A). Therefore, when computing the optimal number of clusters for PathBank pathways (Figure 5A) we automated the silhouette score procedure by smoothing the z-score curve and selecting the minimum value of *k* for which the z-score dropped below 70% of its maximum value, as the optimal number of clusters. We then computed a z-score for the observed number of clusters in the real pathway from these distributions. Since pathways have different numbers of genes, we generated a distinct null distribution for each pathway using the same number of genes as in the pathway itself (Figure 5-figure supplement 1B). Finally, we ranked the pathways based on their deviation from this matched null distribution. Some pathways showed signatures of recurrence (lower number of clusters than expected), whereas others showed more clusters than expected (an indication of high specificity across cell states) (Figure 5B). Additionally, we computed a p-value for each pathway based on the fraction of random sets of genes with higher deviation. This p-value allowed us to identify the most significant pathways (Figure 5B - blue dots). However, we notice that an empirical p-value might be sensitive to the estimation of the null distribution and therefore decided to focus on the rank to identify the top recurrent and cell-state specific pathways.

### Interpathway correlations

To detect potential statistical dependence between pathway states from different signaling pathways, we computed a pairwise Adjusted Mutual Information (AMI) for each pair of pathways. The AMI quantifies statistical dependencies between categorical features in a dataset. In this case, each cell state has two different categorical labels, one for each pathway. The AMI accounts for the expected correlations if the two labels are assigned at random. An AMI value of 0 represents the expected co-occurrence of labels due to chance, while a value of 1 represents perfect statistical dependence between the two clusterings.

### Pseudotime trajectory analysis on developmental datasets

To study transitions in pathway signaling profiles through the course of developmental processes, we performed pseudotime trajectory analysis on two developmental datasets that were not included in the main integrated data set (Figure 2): the neural crest developmental lineage from embryonic day 9.5 (Soldatov et al. 2019), and the haemato-endothelial lineages from embryonic development days 7.5 to 8.5 subsetted from a scRNA-seq atlas of early organogenesis (Pijuan-Sala et al., 2019). We clustered single-cell data as described above (*Clustering single cells and defining cell states*) and constructed a force-directed projection using the ForceAtlas2 algorithm (Jacomy, 2011). We used cluster annotations and the ForceAtlas2 reduced dimensional space as input to the Slingshot algorithm (Street et al., 2018) to obtain a global lineage structure. We then placed cell states in the ordering given by the resulting pseudotime coordinates (Figure 6 A, C). For comparison with integrated atlas counts, the counts from these developmental datasets were scaled in a similar manner to the integrated atlas (Figure 6 B,D). Finally, we used the k-nearest neighbors algorithm to obtain the profile numbers which match a given cell state along a developmental trajectory (Figure 6 B, D, numbers).

---

**Figure 2, Supplement 1.**
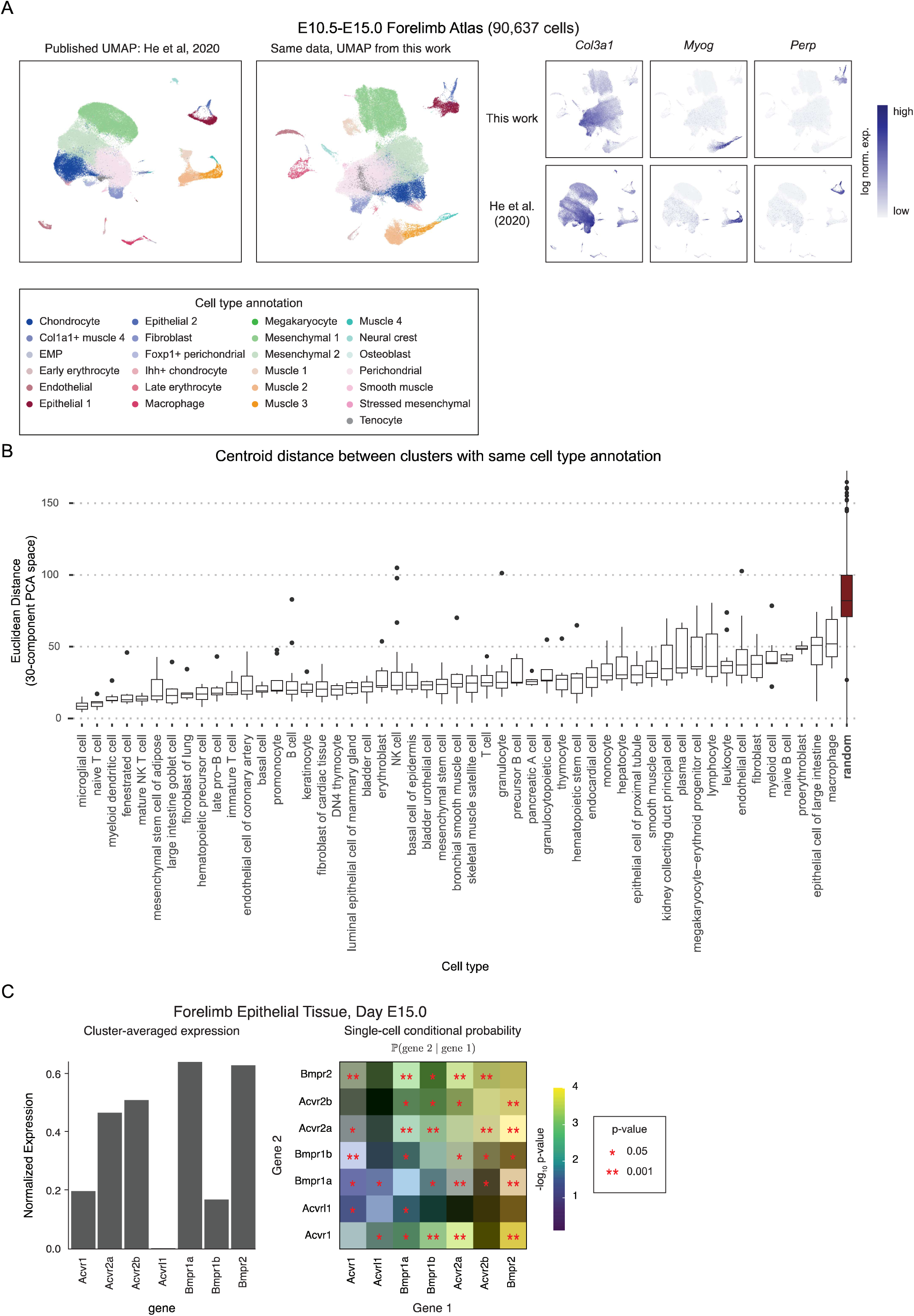
A. Analysis of scRNA-seq datasets using the standard Scanpy pipeline recapitulates published analyses, including (He et al., 2020). Independent analysis of mouse forelimb over days E10.5-E15.0 shows similar cell types (colors, left) and gene expression (right). B. The integrated atlas captures cell type similarity across datasets. Cell clusters with similar annotations in different data sets remain similar to each other in the integrated atlas. C. Cluster-averaged profiles reflect co-expression in single-cells. Shown is an example of a single cluster from the forelimb epithelial tissue data set at day E15. Left, expression of TGF-β receptor genes averaged over all cells in the cluster corresponding to forelimb epithelial tissue at day E15.0. Right, pairwise conditional probability in single cells of gene 2 expression conditioned on gene 1 expression. Pairs of genes with significant entries (**) are co-expressed in the cluster-averaged profile. Higher-order conditional probabilities were not computed due to dropout effects in scRNA-seq data.

**Figure 3, Supplement 1.**
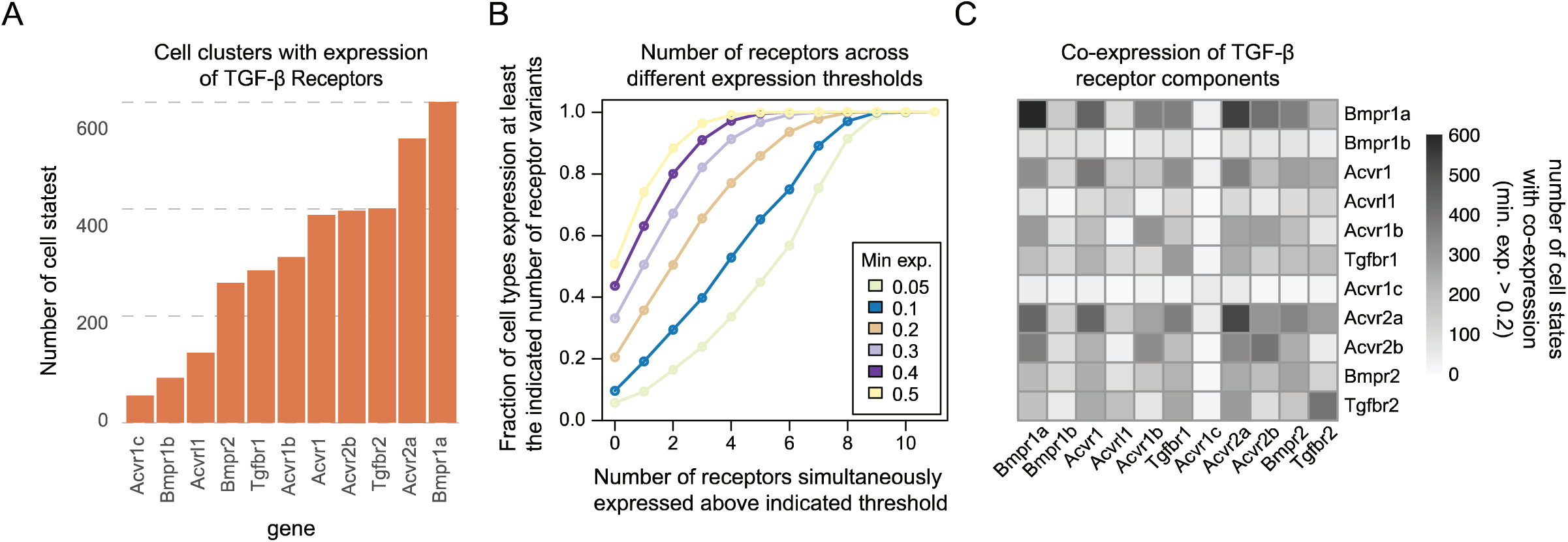
Further analysis of TGF-β pathway expression profiles. A. Histogram showing the number of cell types in the integrated atlas with normalized expression of TGF-β receptors above a threshold of 0.2 in standardized expression units. B. Number of TGF-β receptor components simultaneously expressed for different values of the minimum expression threshold (colors). C. Pairwise co-expression of TGF-β receptor expression reveals broad receptor co-expression patterns. Off-diagonal elements indicate the number of cell states co-expressing, above threshold, the indicated pair of components. Diagonal elements indicate the number of cell states expressing the corresponding individual gene.

**Figure 3, Supplement 2.**
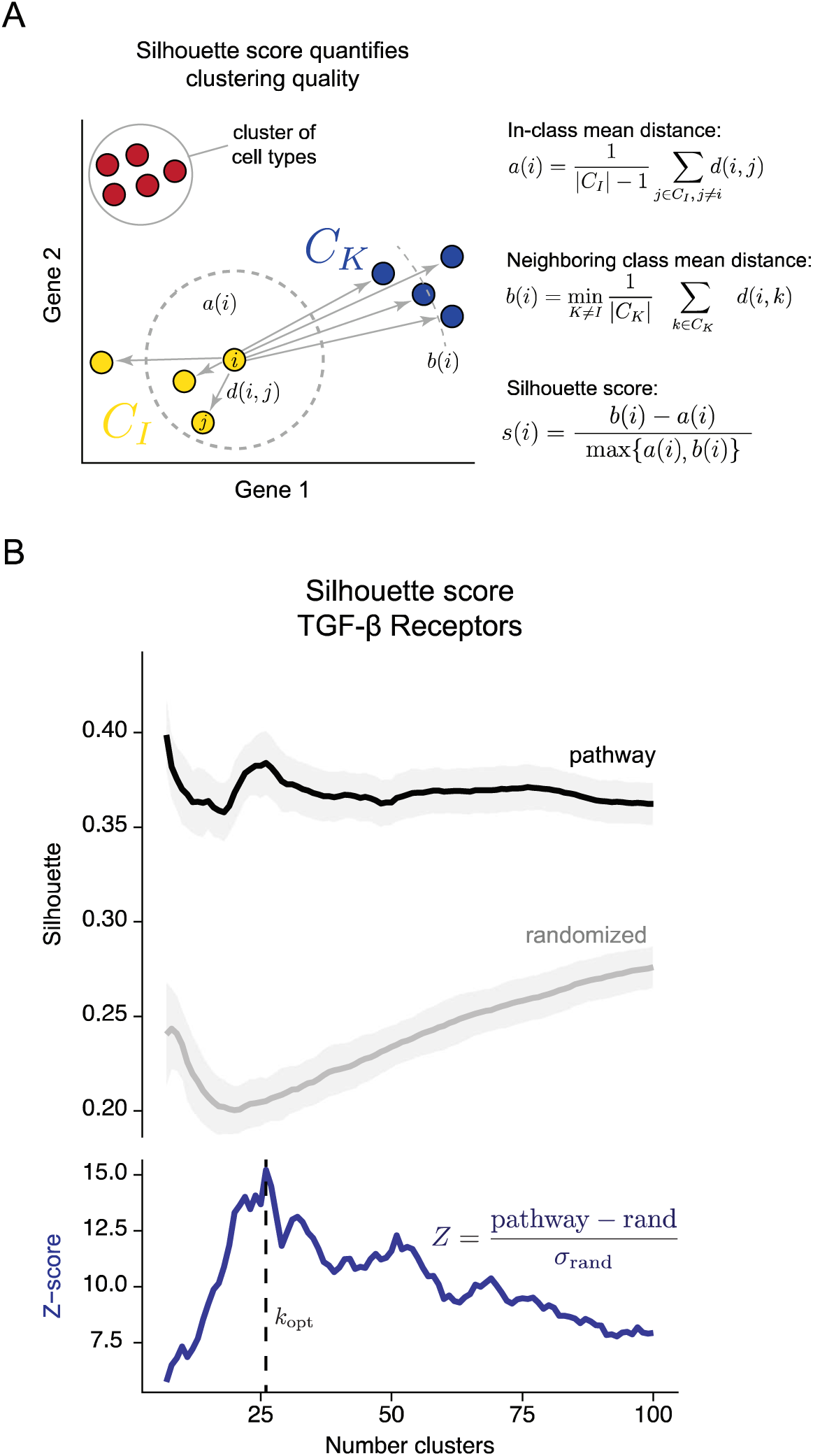
Silhouette analysis can be used to identify optimal clustering thresholds. A. The silhouette score quantifies clustering quality (schematic). For a given clustering, we compute the silhouette score on every data point *i*. We compute *a*(*i*), the mean distance between *i* and every other point in the same cluster, and *b*(*i*), the mean distance between *i* and the nearest neighboring cluster. The silhouette score for data point *i* is then defined as the difference between the inter- and intra-cluster distances, normalized to the maximum of the two (equations). A silhouette score value close to 1 corresponds to well-defined clusters, where data point *i* is similar to other members of its cluster and dissimilar to other clusters, while a value close to -1 suggests poor cluster assignment. The silhouette score for a given clustering, is taken as the average of the individual scores for all data points. B. The silhouette score identifies the approximate number of unique TGF-β receptor expression profiles. We computed the silhouette score across expression values of the pathway genes (black), as well as for 100 random gene sets (gray) where pathway gene expression was independently scrambled for each gene. We then computed the z-score (blue), defined as the silhouette score for pathway genes normalized to the silhouette score for randomized gene sets. We defined the optimal number of receptor profiles *k_opt_* as the number of clusters that produced the peak z-score value (dashed line).

**Figure 4, Supplement 1.**
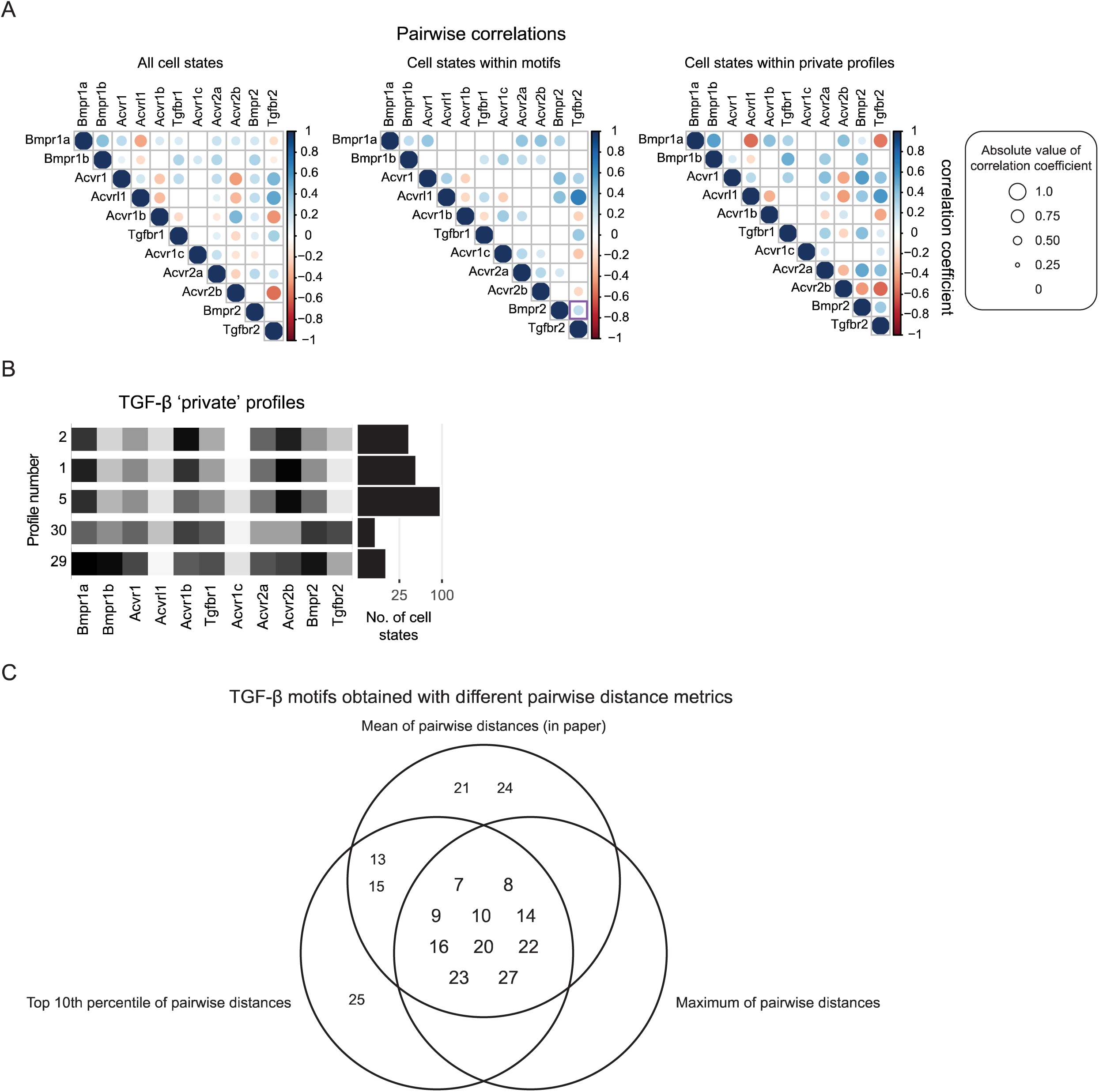
Pairwise correlations among TGF-β receptors and identification of private profiles. A. TGF-β profiles exhibit unique pairwise receptor correlations. Each matrix represents the correlation coefficient for each pair of receptors across all cell states (left), cell states associated with motifs (middle), cell states associated with private profiles (right). B. TGF-β profiles with less than 30 percentile cell type dispersion were classified as *private profiles*. We identified 5 such profiles for TGF-β. Profiles 1, 2, and 5 come from developmental states, while 29 and 30 represent adult cell types. C. Alternative definitions of the dispersion metric recover similar sets of motifs. The mean of intra-class pairwise distances was used as the dispersion metric throughout this work, but we tested two additional dispersion metrics, one that uses the maximum of intra-class pairwise distances, the second that uses the top 10th percentile. The Venn diagram shows profiles identified as motifs from these three distinct definitions of the dispersion metric. The majority of profiles (shown in the intersection of the three circles) are robust to the definition of dispersion. Notably, the dispersion metric that utilizes the maximum of pairwise distances only captures profiles in this intersection. The mean pairwise distance, however, captures two additional profiles as motifs, profiles 21 and 24. Profile 24 contains only two cell states, liver B cells and bone marrow NK cells. The top 10th percentile of pairwise distances captures the adult endothelium-specific profile, 25, as a motif. However, the maximum metric omits profiles 13 and 15, even though they appear to be motifs, since they are both dispersed across the adult smooth muscle and adult kidney epithelium.

**Figure 5, Supplement 1.**
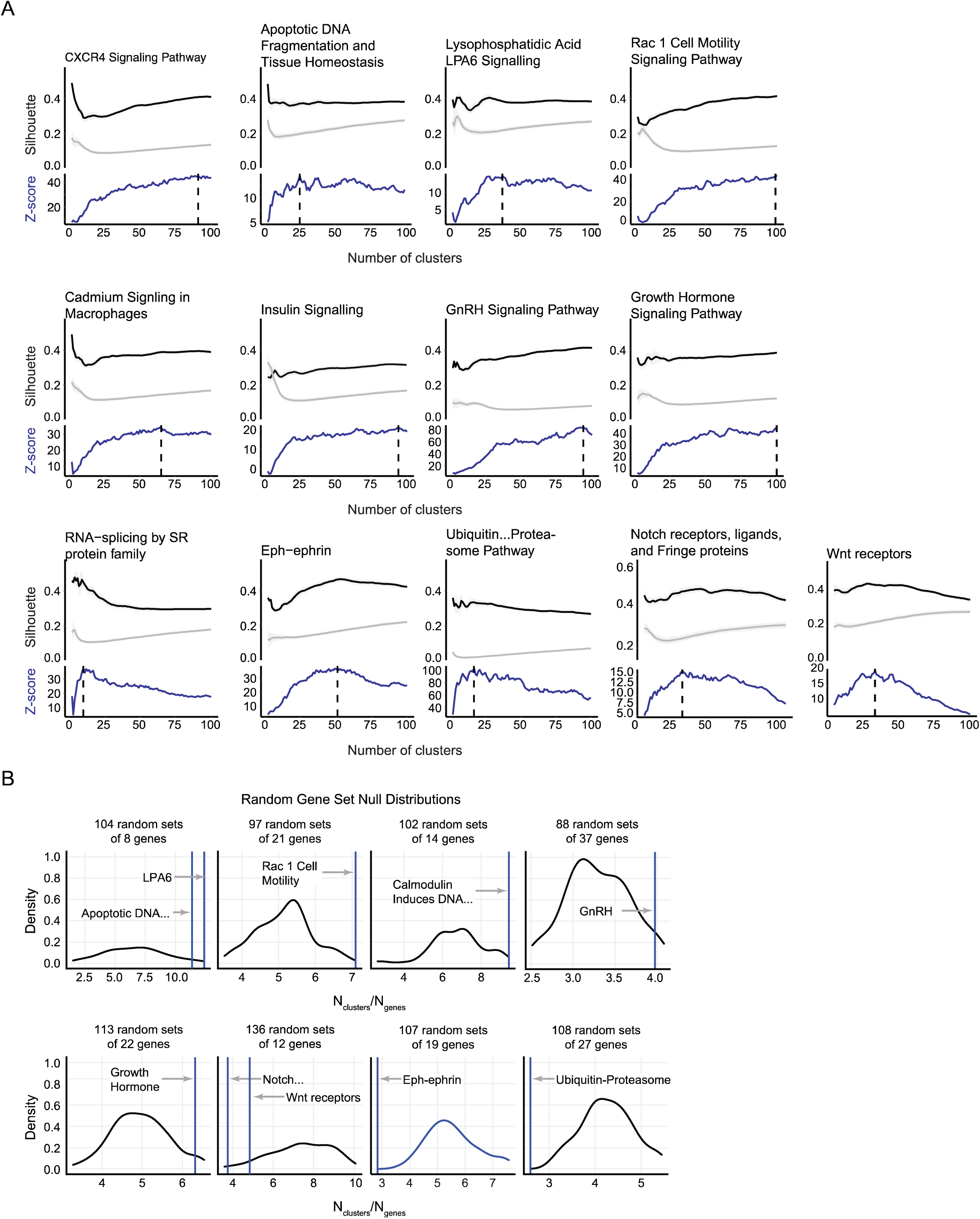
Silhouette profiles for various pathways. A. Silhouette analysis of indicated pathways, as in Figure 3-figure supplement 2B. B. Gene set null distributions for various pathways, as in Figure 5A.

**Figure 5, Supplement 2.**
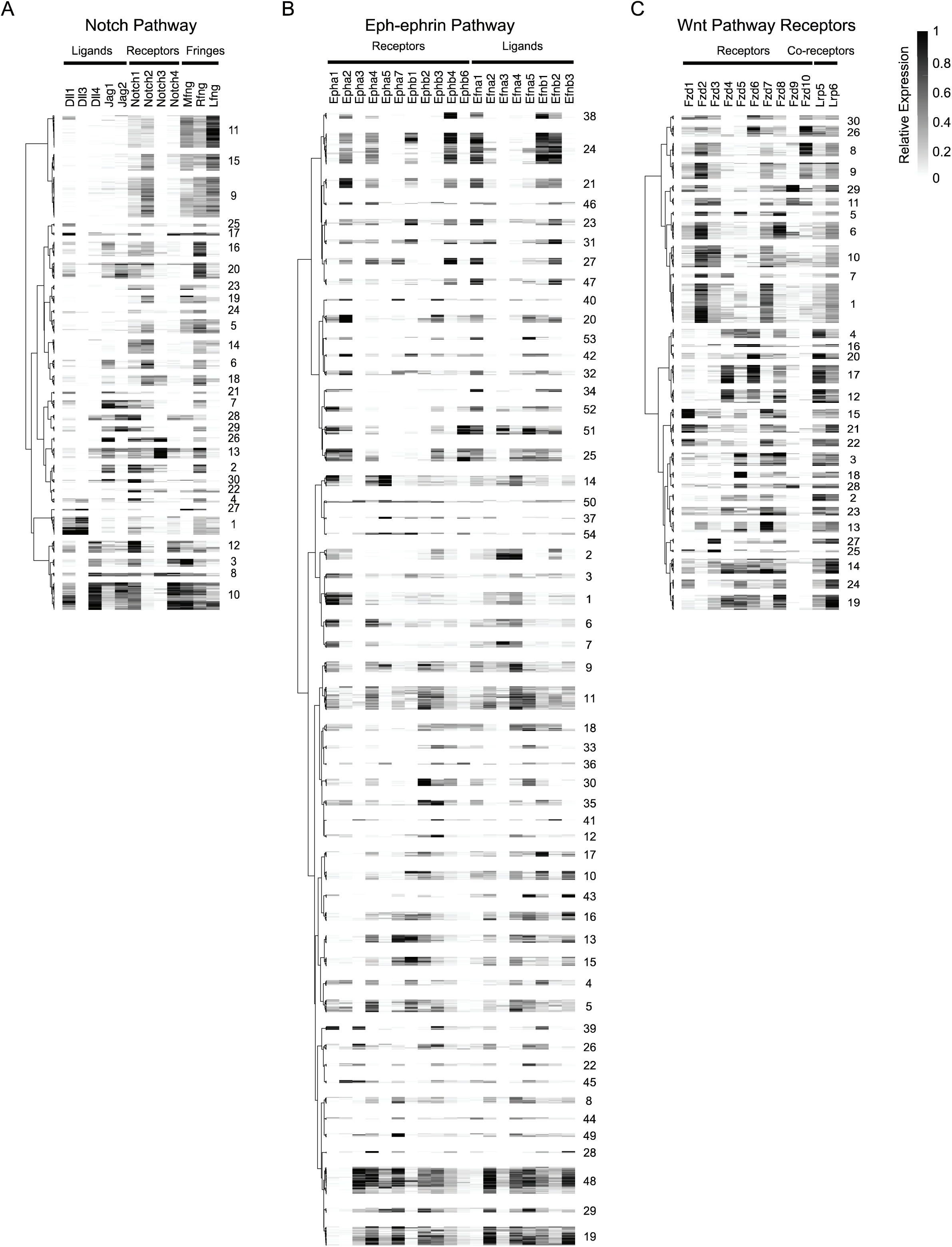
Pathway profiles for Notch, Eph-ephrin, and Wnt receptor receptors. A-C. For each pathway, all pathway profiles are indicated with corresponding labels, as in Figure 2A.

**Figure 5, Supplement 3.**
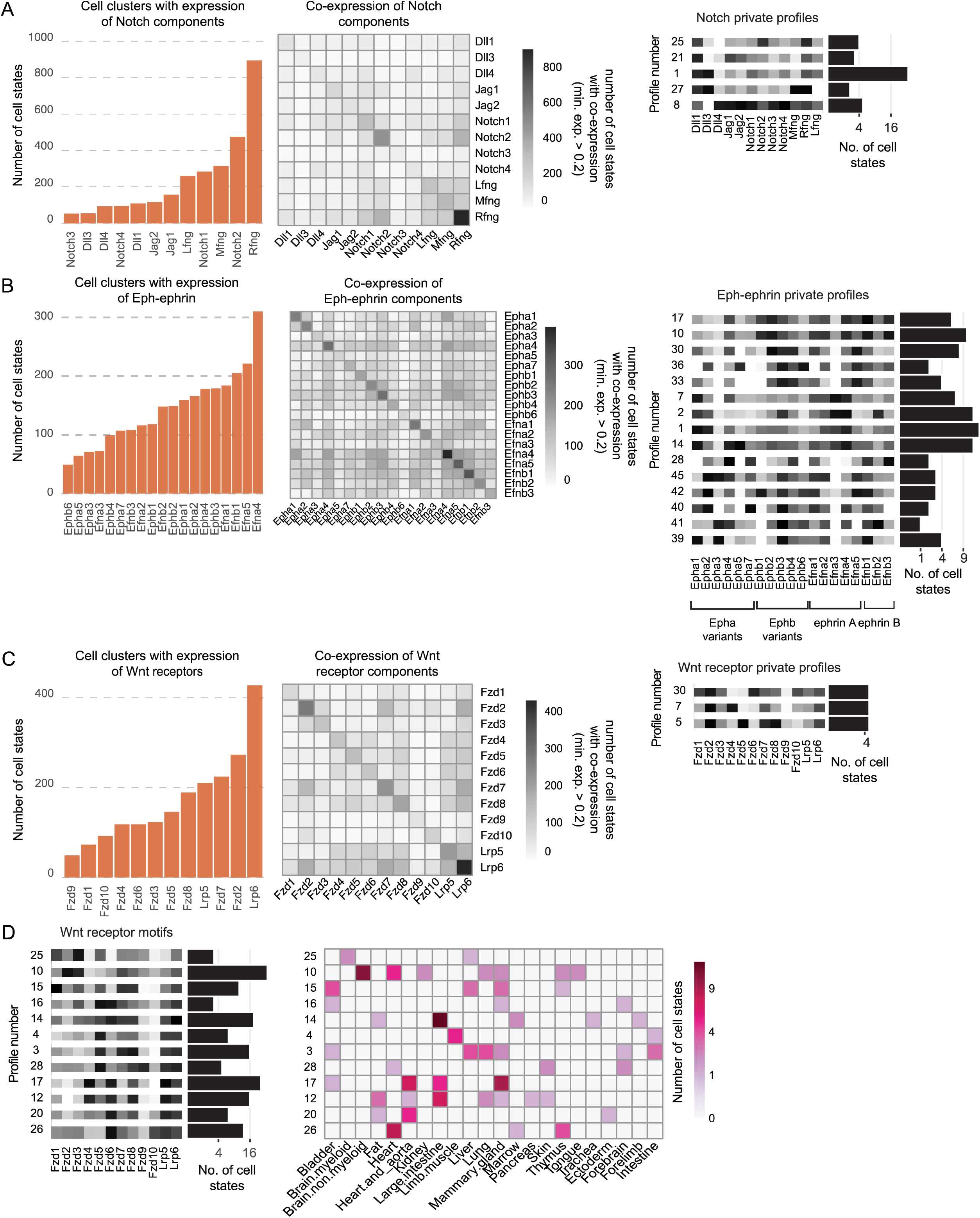
Pathway component prevalence and private profiles for Notch, Eph-ephrin, and Wnt pathways. A-C. Left: Histogram showing the number of cell types in the integrated atlas with normalized expression of Notch (A), Eph-ephrin (B), or Wnt (C) components above a threshold of 0.2 in standardized expression units. Center: Pairwise co-expression analysis of indicated pathway components. Off-diagonal elements indicate the number of cell states co-expressing, above threshold, the indicated pair of components. Diagonal elements indicate the number of cell states expressing the corresponding individual gene. Right: Private profiles for each pathway. Each profile is shown alongside the number of cell states in which it appears (histogram, far right). D. Wnt pathway motifs and their distribution across tissues and organs. These plots are similar to Figure 4D,E and 5C,D but for the Wnt pathway.

## Notes

### Competing Interest Statement

The authors have declared no competing interest.

## References

Antebi, Y. E., Linton, J. M., Klumpe, H., Bintu, B., Gong, M., Su, C., McCardell, R., & Elowitz, M. B. (2017). Combinatorial Signal Perception in the BMP Pathway. Cell, 170(6), 1184–1196.e24.

Antebi, Y. E., Nandagopal, N., & Elowitz, M. B. (2017). An operational view of intercellular signaling pathways. Current Opinion in Systems Biology, 1, 16–24.

Artavanis-Tsakonas, S., Rand, M. D., & Lake, R. J. (1999). Notch signaling: cell fate control and signal integration in development. Science, 284(5415), 770–776.

Arthur, A., & Gronthos, S. (2021). Eph-Ephrin Signaling Mediates Cross-Talk Within the Bone Microenvironment. Frontiers in Cell and Developmental Biology, 9, 598612.

Astin, J. W., Batson, J., Kadir, S., Charlet, J., Persad, R. A., Gillatt, D., Oxley, J. D., & Nobes, C. D. (2010). Competition amongst Eph receptors regulates contact inhibition of locomotion and invasiveness in prostate cancer cells. Nature Cell Biology, 12(12), 1194–1204.

Bhatt, S., Diaz, R., & Trainor, P. A. (2013). Signals and switches in Mammalian neural crest cell differentiation. Cold Spring Harbor Perspectives in Biology, 5(2). https://doi.org/10.1101/cshperspect.a008326

Buckles, G. R., Thorpe, C. J., Ramel, M.-C., & Lekven, A. C. (2004). Combinatorial Wnt control of zebrafish midbrain-hindbrain boundary formation. Mechanisms of Development, 121(5), 437–447.

Canu, G., & Ruhrberg, C. (2021). First blood: the endothelial origins of hematopoietic progenitors. Angiogenesis. https://doi.org/10.1007/s10456-021-09783-9

Chen, H., Brady Ridgway, J., Sai, T., Lai, J., Warming, S., Chen, H., Roose-Girma, M., Zhang, G., Shou, W., & Yan, M. (2013). Context-dependent signaling defines roles of BMP9 and BMP10 in embryonic and postnatal development. Proceedings of the National Academy of Sciences of the United States of America, 110(29), 11887–11892.

Cramer, K. S., & Miko, I. J. (2016). Eph-ephrin signaling in nervous system development. F1000Research, 5. https://doi.org/10.12688/f1000research.7417.1

David, C. J., & Massagué, J. (2018). Contextual determinants of TGFβ action in development, immunity and cancer. Nature Reviews. Molecular Cell Biology, 19(7), 419–435.

del Álamo, D., Rouault, H., & Schweisguth, F. (2011). Mechanism and significance of cis-inhibition in Notch signalling. Current Biology: CB, 21(1), R40–R47.

Derynck, R., & Budi, E. H. (2019). Specificity, versatility, and control of TGF-β family signaling. Science Signaling, 12(570). https://doi.org/10.1126/scisignal.aav5183

Dong, J., Hu, Y., Fan, X., Wu, X., Mao, Y., Hu, B., Guo, H., Wen, L., & Tang, F. (2018). Single-cell RNA-seq analysis unveils a prevalent epithelial/mesenchymal hybrid state during mouse organogenesis. Genome Biology, 19(1), 31.

D’Souza, B., Miyamoto, A., & Weinmaster, G. (2008). The many facets of Notch ligands. Oncogene, 27(38), 5148–5167.

Dudanova, I., & Klein, R. (2011). The axon’s balancing act: cis- and trans-interactions between Ephs and ephrins [Review of *The axon’s balancing act: cis- and trans-interactions between Ephs and ephrins*]. Neuron, 71(1), 1–3.

Eubelen, M., Bostaille, N., Cabochette, P., Gauquier, A., Tebabi, P., Dumitru, A. C., Koehler, M., Gut, P., Alsteens, D., Stainier, D. Y. R., Garcia-Pino, A., & Vanhollebeke, B. (2018). A molecular mechanism for Wnt ligand-specific signaling. Science, 361(6403). https://doi.org/10.1126/science.aat1178

Gerhart, J. (1999). 1998 Warkany lecture: signaling pathways in development. Teratology, 60(4), 226–239.

Goentoro, L., & Kirschner, M. W. (2009). Evidence that fold-change, and not absolute level, of beta-catenin dictates Wnt signaling. Molecular Cell, 36(5), 872–884.

Grigoryan, T., Wend, P., Klaus, A., & Birchmeier, W. (2008). Deciphering the function of canonical Wnt signals in development and disease: conditional loss- and gain-of-function mutations of beta-catenin in mice. Genes & Development, 22(17), 2308–2341.

Groot, A. J., Habets, R., Yahyanejad, S., Hodin, C. M., Reiss, K., Saftig, P., Theys, J., & Vooijs, M. (2014). Regulated proteolysis of NOTCH2 and NOTCH3 receptors by ADAM10 and presenilins. Molecular and Cellular Biology, 34(15), 2822–2832.

Grosswendt, S., Kretzmer, H., Smith, Z. D., Kumar, A. S., Hetzel, S., Wittler, L., Klages, S., Timmermann, B., Mukherji, S., & Meissner, A. (2020). Epigenetic regulator function through mouse gastrulation. Nature, 584(7819), 102–108.

He, P., Williams, B. A., Trout, D., Marinov, G. K., Amrhein, H., Berghella, L., Goh, S.-T., Plajzer-Frick, I., Afzal, V., Pennacchio, L. A., Dickel, D. E., Visel, A., Ren, B., Hardison, R. C., Zhang, Y., & Wold, B. J. (2020). The changing mouse embryo transcriptome at whole tissue and single-cell resolution. Nature, 583(7818), 760–767.

Jacomy, M. (2011). Force atlas 2 layout.

Kageyama, R., Shimojo, H., & Isomura, A. (2018). Oscillatory Control of Notch Signaling in Development. Advances in Experimental Medicine and Biology, 1066, 265–277.

Kakuda, S., & Haltiwanger, R. S. (2017). Deciphering the Fringe-Mediated Notch Code: Identification of Activating and Inhibiting Sites Allowing Discrimination between Ligands. Developmental Cell, 40(2), 193–201.

Kakuda, S., LoPilato, R. K., Ito, A., & Haltiwanger, R. S. (2020). Canonical Notch ligands and Fringes have distinct effects on NOTCH1 and NOTCH2. The Journal of Biological Chemistry, 295(43), 14710–14722.

Kania, A., & Klein, R. (2016). Mechanisms of ephrin–Eph signalling in development, physiology and disease. Nature Reviews. Molecular Cell Biology, 17(4), 240–256.

Kléber, M., Lee, H.-Y., Wurdak, H., Buchstaller, J., Riccomagno, M. M., Ittner, L. M., Suter, U., Epstein, D. J., & Sommer, L. (2005). Neural crest stem cell maintenance by combinatorial Wnt and BMP signaling. The Journal of Cell Biology, 169(2), 309–320.

Klein, R. (2012). Eph/ephrin signalling during development. Development, 139(22), 4105–4109.

Klumpe, H., Langley, M. A., Linton, J. M., Su, C. J., Antebi, Y. E., & Elowitz, M. B. (2020). The context-dependent, combinatorial logic of BMP signaling. In bioRxiv (p. 2020.12.08.416503). https://doi.org/10.1101/2020.12.08.416503

Lafkas, D., Shelton, A., Chiu, C., de Leon Boenig, G., Chen, Y., Stawicki, S. S., Siltanen, C., Reichelt, M., Zhou, M., Wu, X., Eastham-Anderson, J., Moore, H., Roose-Girma, M., Chinn, Y., Hang, J. Q., Warming, S., Egen, J., Lee, W. P., Austin, C., … Siebel, C. W. (2015). Therapeutic antibodies reveal Notch control of transdifferentiation in the adult lung. Nature, 528(7580), 127–131.

LeBon, L., Lee, T. V., Sprinzak, D., Jafar-Nejad, H., & Elowitz, M. B. (2014). Fringe proteins modulate Notch-ligand cis and trans interactions to specify signaling states. eLife, 3, e02950.

Lim, W., Mayer, B., & Pawson, T. (2015). Cell Signaling: Principles and Mechanisms. Garland Science.

Li, P., & Elowitz, M. B. (2019). Communication codes in developmental signaling pathways. Development, 146(12). https://doi.org/10.1242/dev.170977

MacDonald, B. T., & He, X. (2012). Frizzled and LRP5/6 receptors for Wnt/β-catenin signaling. Cold Spring Harbor Perspectives in Biology, 4(12). https://doi.org/10.1101/cshperspect.a007880

Massagué, J. (2012). TGFβ signalling in context. Nature Reviews. Molecular Cell Biology, 13(10), 616–630.

Merlos-Suárez, A., & Batlle, E. (2008). Eph–ephrin signalling in adult tissues and cancer. Current Opinion in Cell Biology, 20(2), 194–200.

Mikels, A. J., & Nusse, R. (2006). Wnts as ligands: processing, secretion and reception. Oncogene, 25(57), 7461–7468.

Muñoz Descalzo, S., & Martinez Arias, A. (2012). The structure of Wntch signaling and the resolution of transition states in development. Seminars in Cell & Developmental Biology, 23(4), 443–449.

Nandagopal, N., Santat, L. A., & Elowitz, M. B. (2019). activation in the Notch signaling pathway. eLife, 8. https://doi.org/10.7554/eLife.37880

Nowotschin, S., Setty, M., Kuo, Y.-Y., Liu, V., Garg, V., Sharma, R., Simon, C. S., Saiz, N., Gardner, R., Boutet, S. C., Church, D. M., Hoodless, P. A., Hadjantonakis, A.-K., & Pe’er, D. (2019). The emergent landscape of the mouse gut endoderm at single-cell resolution. Nature, 569(7756), 361–367.

Okigawa, S., Mizoguchi, T., Okano, M., Tanaka, H., Isoda, M., Jiang, Y.-J., Suster, M., Higashijima, S.-I., Kawakami, K., & Itoh, M. (2014). Different combinations of Notch ligands and receptors regulate V2 interneuron progenitor proliferation and V2a/V2b cell fate determination. Developmental Biology, 391(2), 196–206.

Pijuan-Sala, B., Griffiths, J. A., Guibentif, C., Hiscock, T. W., Jawaid, W., Calero-Nieto, F. J., Mulas, C., Ibarra-Soria, X., Tyser, R. C. V., Ho, D. L. L., Reik, W., Srinivas, S., Simons, B. D., Nichols, J., Marioni, J. C., & Göttgens, B. (2019a). A single-cell molecular map of mouse gastrulation and early organogenesis. Nature, 566(7745), 490–495.

Pijuan-Sala, B., Griffiths, J. A., Guibentif, C., Hiscock, T. W., Jawaid, W., Calero-Nieto, F. J., Mulas, C., Ibarra-Soria, X., Tyser, R. C. V., Ho, D. L. L., Reik, W., Srinivas, S., Simons, B. D., Nichols, J., Marioni, J. C., & Göttgens, B. (2019b). A single-cell molecular map of mouse gastrulation and early organogenesis. Nature, 566(7745), 490–495.

Qiu, C., Cao, J., Li, T., Srivatsan, S., Huang, X., Calderon, D., Noble, W. S., Disteche, C. M., Spielmann, M., Moens, C. B., Trapnell, C., & Shendure, J. (2021). Systematic reconstruction of the cellular trajectories of mammalian embryogenesis. In bioRxiv (p. 2021.06.08.447626). https://doi.org/10.1101/2021.06.08.447626

Rohani, N., Parmeggiani, A., Winklbauer, R., & Fagotto, F. (2014). Variable combinations of specific ephrin ligand/Eph receptor pairs control embryonic tissue separation. PLoS Biology, 12(9), e1001955.

Rousseeuw, P. J. (1987). Silhouettes: A graphical aid to the interpretation and validation of cluster analysis. In Journal of Computational and Applied Mathematics (Vol. 20, pp. 53–65). https://doi.org/10.1016/0377-0427(87)90125-7

Salvucci, O., & Tosato, G. (2012). Essential roles of EphB receptors and EphrinB ligands in endothelial cell function and angiogenesis. Advances in Cancer Research, 114, 21–57.

Seiradake, E., Schaupp, A., del Toro Ruiz, D., Kaufmann, R., Mitakidis, N., Harlos, K., Aricescu, R., Klein, R., & Jones, E. Y. (2013). Structurally encoded intraclass differences in EphA clusters drive distinct cell responses. Nature Structural & Molecular Biology, 20(8), 958–964.

Siebel, C., & Lendahl, U. (2017). Notch Signaling in Development, Tissue Homeostasis, and Disease. Physiological Reviews, 97(4), 1235–1294.

Simões-Costa, M., & Bronner, M. E. (2015). Establishing neural crest identity: a gene regulatory recipe. Development, 142(2), 242–257.

Soldatov, R., Kaucka, M., Kastriti, M. E., Petersen, J., Chontorotzea, T., Englmaier, L., Akkuratova, N., Yang, Y., Häring, M., Dyachuk, V., Bock, C., Farlik, M., Piacentino, M. L., Boismoreau, F., Hilscher, M. M., Yokota, C., Qian, X., Nilsson, M., Bronner, M. E., … Adameyko, I. (2019). Spatiotemporal structure of cell fate decisions in murine neural crest. Science, 364(6444). https://doi.org/10.1126/science.aas9536

Sprinzak, D., Lakhanpal, A., Lebon, L., Santat, L. A., Fontes, M. E., Anderson, G. A., Garcia-Ojalvo, J., & Elowitz, M. B. (2010). Cis-interactions between Notch and Delta generate mutually exclusive signalling states. Nature, 465(7294), 86–90.

Street, K., Risso, D., Fletcher, R. B., Das, D., Ngai, J., Yosef, N., Purdom, E., & Dudoit, S. (2018). Slingshot: cell lineage and pseudotime inference for single-cell transcriptomics. BMC Genomics, 19(1), 477.

Su, C. J., Murugan, A., Linton, J. M., Yeluri, A., Bois, J., Klumpe, H., Antebi, Y. E., & Elowitz, M. (2020). Ligand-receptor promiscuity enables cellular addressing. In bioRxiv (p. 2020.12.08.412643). https://doi.org/10.1101/2020.12.08.412643

Tabula Muris Consortium. (2020). A single-cell transcriptomic atlas characterizes ageing tissues in the mouse. Nature, 583(7817), 590–595.

Tabula Muris Consortium, Overall coordination, Logistical coordination, Organ collection and processing, Library preparation and sequencing, Computational data analysis, Cell type annotation, Writing group, Supplemental text writing group, & Principal investigators. (2018). Single-cell transcriptomics of 20 mouse organs creates a Tabula Muris. Nature, 562(7727), 367–372.

Traag, V. A., Waltman, L., & van Eck, N. J. (2019). From Louvain to Leiden: guaranteeing well-connected communities. Scientific Reports, 9(1), 5233.

Tual-Chalot, S., Mahmoud, M., Allinson, K. R., Redgrave, R. E., Zhai, Z., Oh, S. P., Fruttiger, M., & Arthur, H. M. (2014). Endothelial depletion of Acvrl1 in mice leads to arteriovenous malformations associated with reduced endoglin expression. PloS One, 9(6), e98646.

Verkaar, F., & Zaman, G. J. R. (2010). A model for signaling specificity of Wnt/Frizzled combinations through co-receptor recruitment. In FEBS Letters (Vol. 584, Issue 18, pp. 3850–3854). https://doi.org/10.1016/j.febslet.2010.08.030

Vilar, J. M. G., Jansen, R., & Sander, C. (2006). Signal processing in the TGF-beta superfamily ligand-receptor network. PLoS Computational Biology, 2(1), e3.

Voloshanenko, O., Gmach, P., Winter, J., Kranz, D., & Boutros, M. (2017). Mapping of Wnt-Frizzled interactions by multiplex CRISPR targeting of receptor gene families. FASEB Journal: Official Publication of the Federation of American Societies for Experimental Biology, 31(11), 4832–4844.

Wang, Y., Chang, H., Rattner, A., & Nathans, J. (2016). Frizzled Receptors in Development and Disease. Current Topics in Developmental Biology, 117, 113–139.

Wishart, D. S., Li, C., Marcu, A., Badran, H., Pon, A., Budinski, Z., Patron, J., Lipton, D., Cao, X., Oler, E., Li, K., Paccoud, M., Hong, C., Guo, A. C., Chan, C., Wei, W., & Ramirez-Gaona, M. (2020). PathBank: a comprehensive pathway database for model organisms. Nucleic Acids Research, 48(D1), D470–D478.

Wolf, F. A., Angerer, P., & Theis, F. J. (2018). SCANPY: large-scale single-cell gene expression data analysis. Genome Biology, 19(1), 15.

Wrana, J. L., Attisano, L., Cárcamo, J., Zentella, A., Doody, J., Laiho, M., Wang, X.-F., & Massague, J. (1992). TGFβ signals through a heteromeric protein kinase receptor complex. In Cell (Vol. 71, Issue 6, pp. 1003–1014). https://doi.org/10.1016/0092-8674(92)90395-s

Wurdak, H. (2005). Inactivation of TGF signaling in neural crest stem cells leads to multiple defects reminiscent of DiGeorge syndrome. In Genes & Development (Vol. 19, Issue 5, pp. 530–535). https://doi.org/10.1101/gad.31740

